# Enabling Transparent Toxicokinetic Modeling for Public Health Risk Assessment

**DOI:** 10.1101/2024.08.19.608571

**Authors:** Sarah E. Davidson-Fritz, Caroline L. Ring, Marina V. Evans, Celia M. Schacht, Xiaoqing Chang, Miyuki Breen, Gregory S. Honda, Elaina Kenyon, Matthew W. Linakis, Annabel Meade, Robert G. Pearce, Mark A. Sfeir, James P. Sluka, Michael J. Devito, John F. Wambaugh

**Author notes:** Author Contributions: SEF: Conceptualization, Data Curation, Investigation, Project Administration, Software, Supervision, Visualization, Writing – Original Draft Preparation CLR: Investigation, Methodology, Software, Validation MVE: Methodology, Writing – Review & Editing CMS: Investigation, Methodology, Software XC: Investigation, Methodology MB: Investigation, Software GSH: Conceptualization, Methodology, Software EK: Validation, Writing – Review & Editing MWL: Conceptualization, Methodology, Software, Writing – Review & Editing AM: Investigation, Methodology, Software RGP: Conceptualization, Data Curation, Investigation, Methodology, Software, Validation MAS: Conceptualization, Data Curation, Software JPS: Investigation, Writing – Review & Editing MJD: Conceptualization, Funding Acquisition, Resources, Writing – Review & Editing JFW: Conceptualization, Data Curation, Funding Acquisition, Investigation, Methodology, Project Administration, Resources, Software, Supervision, Validation, Visualization, Writing – Original Draft Preparation.

## Abstract

Toxicokinetics describes the absorption, distribution, metabolism, and elimination of chemicals by the body. Predictions from toxicokinetic models provide key information for chemical risk assessment. Traditionally, these predictions extrapolate from experimental animal species data (for example, in rats) to humans. More recently, toxicokinetics has been used for extrapolation from *in vitro* “new approach methods (NAMs)” for toxicology to *in vivo*. Chemical-specific *in vivo* toxicokinetic data are often unavailable for the thousands of chemicals in commerce. Therefore, large amounts of *in vitro* data measuring chemical-specific toxicokinetics have been collected. These data enable “high-throughput toxicokinetic” or HTTK modeling. The *httk* R package provides a library of chemical-specific data from peer-reviewed HTTK studies. *httk* further provides a suite of tools for parameterizing and evaluating toxicokinetic models. *httk* uses the open-source language MCSim to describe models for compartmental and physiologically based toxicokinetics (PBTK), MCSim can convert the model descriptions into a high-speed C code script. New models are integrated into *httk* using the open-source package development functionality in R, a model documentation file (R script), and the HTTK model description code file (C script). In addition to HTTK models, *httk* provides a series of functionalities such as unit conversion, model parameterization, Monte Carlo simulations for uncertainty propagation and biological variability, *in vivo*-derived data for evaluating model predictions, and other model utility functions. Here, we describe in detail how to add new HTTK models to *httk* and take advantage of the pre-existing data and functionality in the package. As a demonstration, we describe the integration of the gas inhalation PBTK model into *httk*. Modern modeling approaches, as exemplified by *httk*, allow for clear communication, reproducibility, and public scrutiny. The intention of *httk* is to provide a transparent, open-source tool for toxicokinetics, bioinformatics, and public health risk assessment.

**Author Summary:** We describe the integration and evaluation of new physiologically based toxicokinetic (PBTK) models into an open-source R package. Adding a new model to the R package allows a modeler to use the existing tools and data for *in vitro* to *in vivo* extrapolation (IVIVE). Integration with the R statistical analysis environment further allows model assessment. This workflow is designed to create a more transparent and reproducible approach to toxicokinetic models developed for various exposure scenarios. Here, we demonstrate the model integration and evaluation workflow with an inhalation model. Additionally, we provide an evaluation of the overall package performance as new models, data, and functionality are added over time. Our results show that transparent development of models, and use of existing data within the open-source R package format, allows for improvement of *in vitro* to *in vivo* extrapolation estimations. IVIVE is vital for advancement of 21^st^ century human health risk assessment.

## Introduction

High-throughput toxicokinetics (HTTK) allows the incorporation of chemical-specific toxicokinetic (TK) predictions into bioinformatic workflows analyzing effects across large numbers of chemicals. These workflows in turn enable more rapid chemical risk assessment. Next generation risk assessment relies in part on toxicological *in vitro-in vivo* extrapolation (IVIVE) to relate bioactive concentrations identified *in vitro* to real world doses to which people might actually be exposed [1–4]. In HTTK, chemicals are characterized by standardized *in vitro* measurements, or *in silico* predictions [5]. That is, HTTK uses *toxicokinetic* IVIVE to allow *toxicological* IVIVE for risk assessment. A series of chemical-independent (“generic”) toxicokinetic models have been created for relating chemical-specific values and external chemical exposures to internal tissue doses. The U.S. EPA’s R package *httk* [6] is one example of an HTTK tool originally designed not so much to accommodate development of toxicokinetic models, but rather to facilitate the incorporation of toxicokinetics seamlessly into chemical risk analysis workflows. (Throughout this document we distinguish the general science of HTTK from the specific implementation of the R package *httk* using capitalization and italics.) The *httk* R package provides a suite of empirical and physiologically based toxicokinetic (PBTK) models [6–8] designed for bioinformatics [9–11]. However, the data and software infrastructure of *httk* provide a platform for potential expansion with new models. These new models can simulate important aspects of toxicokinetics currently unavailable in the suite of existing HTTK models. Here, we describe the process for developing and integrating new HTTK models into *httk*.

Schmolke, Thorbek (12) wrote that “empirical approaches are often too limited to inform policy and decision making” and therefore “decision making requires models”. Decisions about the risk chemicals pose to public health involves models for toxicity, exposure, and the dose-response relationship [13]. Toxicokinetics informs the dose-response relationship [14] by describing the absorption, distribution, metabolism, and elimination of one or more chemical dose(s). Mathematical models for toxicokinetics allow predictions of the relationship between external doses to the body and internal tissue concentrations [15]. Within the pharmaceutical industry PBTK models are widely used to better understand drug absorption and disposition [16, 17]. PBTK models provide the backbone for insights into toxicodynamic effects such as drug-drug interactions [16, 17]. Evaluating confidence in a particular TK model can be done via comparison between HTTK model predictions and *in vivo* measurements [18]. The correspondence between predictions and observations allows for the estimation of bias and uncertainty; decision makers may then consider using models to extrapolate to other situations (for example, dose, route, physiology) where data may be unavailable [19]. However, despite the deterministic nature of scientific computing (even including reproducible pseudo-random number generation [20]) it is often difficult to reproduce computational results [21] due in part to lack of standardized means to distributed, document, and evaluate models [22] and discrepancies that emerge between conducting research with a model and the mechanisms available to describe the model and its analysis in a research manuscript [23]. In non-pharmaceutical chemical risk assessments model reproducibility, statistical evaluation [24], and data availability are all barriers to the incorporation of PBTK models into public health risk assessment [25].

It is desirable to assess the risk posed to public health for tens of thousands of non-pharmaceutical chemicals, both in the environment and commerce [26]. For ethical reasons it is typically infeasible to obtain human toxicokinetic data from controlled exposure(s) to these non-therapeutic chemicals. Controlled exposures are where the magnitude, route, and timing of chemical exposure are sufficiently known to allow mathematical modeling. Furthermore, there are limited resources for obtaining similar data in animals, especially considering the very large number of compounds for which a data is needed. Thus, there is a general lack of *in vivo* data for building and evaluating toxicokinetic models for non-pharmaceuticals [27, 28]. In these cases, generic PBTK models offer an alternative path [29–34]. Generic PBTK models describe both a standardized physiology and a set of standardized parameters for describing chemical specificity [5, 29]. The chemical specific parameters can often be obtained by *in vitro* or *in silico* methods [1, 35]. Because there are many TK processes that are important for only a subset of chemical classes, we expect a generic model to have larger uncertainty than a “bespoke” chemical-specific model [36]. However, the same generic model may be used across many chemicals, which offers greater confidence in model implementation and reproducibility, and may also provide a consistent set of predictions across a wide range of chemicals. The key advantage of using generic models is the ability to evaluate the model in the absence of chemical-specific *in vivo* data [37].

Over the past decade, HTTK data has become publicly available for more than a thousand chemicals [5]. Concurrently, public repositories of *in vivo* toxicokinetic observations (that is, chemical concentration in various tissues as a function of dose and time) have become available to allow statistical analysis of the performance of HTTK [38]. While we may not have *in vivo* data for the chemical of interest, a generic model may be evaluated using as much *in vivo* data as possible for other chemicals described using the same *in vitro* and *in silico* predictors. That is, the evaluation of the HTTK model is not dependent upon the presence of data for a specific chemical of interest, but predictive performance can still be evaluated with other chemicals that do have data. Any inaccurate approximations, omissions, or mistakes in the HTTK model should increase the estimated uncertainty when evaluated systematically across chemicals [37]. The generic nature of the HTTK models allow for estimation of bias, uncertainty, and subsequent correlation of residuals with chemical-specific properties. [37, 39] Decision makers can then judge whether the generic model can be used for a chemical that does not have *in vivo* data but for which *in vitro* or *in silico* HTTK values are available.

Pearce, Setzer (6) described the original release of *httk* which comprised a one- and three-compartment empirical TK model along with a simplified PBTK model. Ring, Pearce (40) and Breen, Wambaugh (41) focused on the Monte Carlo human variability simulator (namely, “httk-pop”). Wambaugh, Wetmore (42) discussed how the Monte Carlo approach also allowed propagation of chemical-specific measurement uncertainty. *httk* uses the built-in R documentation functions – these currently provide substantive detail at the level of individual functions and data sets. Here, we discuss how new HTTK models may be created and integrated with the *httk* package to use previously evaluated tools and data alongside new approaches. Once an HTTK model is developed and integrated with a chemical library it becomes usable in toxicokinetic applications ranging from informing regulatory decision makers [10, 43, 44] to systems biology [11, 45, 46]. For example, inclusion of the Linakis et al.[7] inhalation model provided a new HTTK model addressing an exposure route previously not implemented.

We are inviting the biological modeling community to help expand the physiology and chemical domains addressed by *httk*. This work is intended to help new models use the existing HTTK data and functionality. Whenever such a model is developed, evaluated, and peer-reviewed, the EPA is hopeful to add that new model to the suite of HTTK models. While public release is not required, if a new model is included in the publicly released *httk* R package then that model will be widely available to modelers, bioinformaticists, and risk assessors. Ultimately, the most challenging task is not building a new model, but evaluating its performance [47, 48]. New generic TK models should be statistically evaluated with data in *httk,* which includes a growing library of chemicals. When *in vivo* data are available, each new model incorporated into *httk* should include a comparison of the model predictions relative to *in vivo* data. The evaluation allowed by incorporating a model into *httk* provides an estimate of how generalizable the new (or revised) model is expected to be. “The first principle is that you must not fool yourself—and you are the easiest person to fool.” [49].

## Results

As of 2024, 1046 chemicals have had *in vitro* measured values collected from the literature and made available with *httk*. Each of these chemicals has both *in vitro* measured plasma protein binding and intrinsic hepatic clearance that can be used with generic TK models. Some, simpler TK models can be used with just intrinsic clearance and the assumption that plasma protein binding measurements were attempted but failed because the chemical was highly bound (a default fraction of 0.5% unbound is assumed) [1, 6]. Various structure-based models have been used to provide additional, overlapping sets of *in silico* predictions for 8719 [35], 8,573 [50], and 10,452 [51] chemicals. These *in silico* predictions are available via the commands load_sipes2017(), load_pradeep2020(), and load_dawson2021(), respectively.

To date the model development paradigm is to create new scenario specific models to incorporate into the *httk* package rather than develop a single all-encompassing HTTK model. Each new model should be evaluated for accuracy of model predictions against measured data prior to final incorporation into *httk*. Statistics, such as root mean squared log_10_ error (RMSLE), summarize the concordance between model predictions and data obtained from *in vivo* experiments collected by the chemical concentration vs. time database (CvTdb) [38], Model performance statistics are recorded in *httk* with “vignettes” [52]. For R packages in general, vignettes provide “long-form documentation” with extended examples using functions from the package [53]. For each manuscript describing expansion/refinement of the httk approach we aim to provide a vignette for recreating key figures from that research [5]. Because the vignettes can be rerun with each new package re-build (that is, version), the RMSLE and other performance statistics can be recalculated and may result in new values for one or more of the models. Many of the vignettes are computationally intensive to compile. To mitigate computational time for package rebuilds the beginning of our vignettes include an “execute_vignette” argument with a default of FALSE. However, model developers can set it to TRUE to ensure previous results can be recreated. Our goal is to ensure package revisions made for one model have minimal to no effect on the other models, and if unavoidable that it occurs in a predictable and explainable manner. Here we have developed a new function benchmark_httk() to provide a less computationally intensive tool to calculate many of the relevant RMSLEs and other performance metrics (Table 7).

We have evaluated the performance of the *httk* package at two levels. First, we conducted a case study re-evaluating the predictive accuracy for the Linakis et al.[7] inhalation model. This demonstrates the model performance evaluation process for new HTTK scenarios. Second, we evaluated the overall historical package performance on key HTTK calculations. The second case study examines potentially unintended impacts of package development.

### Case Study Example Model

In Linakis et al.[7] a generic gas inhalation PBTK model was added to *httk*. The files required for adding this model are included as supplemental material. The model was developed through combining a pre-existing generic inhalation model of Jongeneelen and Berge (34) with the inhalation model of Clewell III, Gentry (54), which noted that some chemicals may be absorbed into the mucus or otherwise trapped in the upper respiratory tract (URT). Then the steps of Table 1 (described more fully in the Methods) were followed: First, a model description was written in MCSim (Supplemental File “model_gas_pbtk.model”). This file includes state variables, parameters, allometric scaling and differential equations. The model file was translated to a compilable C script with the “mod” function in MCSim (Supplemental File “model_gas_pbtk-raw.c”). The “raw” C file was then edited to include the needed integrations with other models, data, and tools already present in *httk*. MCSim gives default names to the functions in the newly generated C script and these must be renamed to not overlap with other function (Supplemental File “model_gas_pbtk.c”). The final steps for the model code itself were moving the C script into the “httk/src” directory and then modifying the “httk/src/init.c” file to list the newly create functions in the C script.

**Table 1:** Step-by-Step Instructions for Adding a model to the httk modeling suite. See Methods for elaboration. Find a user guide on-line at: https://github.com/USEPA/CompTox-ExpoCast-httk/blob/main/models/HTTK-Models-Guide.pdf. See supplemental materials for examples. It will likely be necessary for Windows users to install rtools: https://cran.r-project.org/bin/windows/Rtools/rtools44/rtools.html

|  | Step Instructions | Comments |
| --- | --- | --- |
| 1 | Describe a TK model (parameters, equations, unit conversions) using the MCSim Model format | Users can also start by writing their own .c file (skipping to Step 3 if doing so), but the MCSim allows declarative rather than procedural modeling and may be more familiar to the PBTK community (for example, users of the advanced continuous simulation language – ACSL [REF]) |
| 2 | Convert MCSim .model file to .c model file | Requires download and installation of MCSim (MCSim_under_R): https://www.gnu.org/software/mcsim/ |
| 2a | Run building_mod.R file once (in MCSim_under_R > my_R_project folder) | This step generates the mod.exe file required to convert the .model file to the .c file |
| 2b | Run one of the script_compile_run.R files (in MCSim_under_R > my_R_project folder) | This step will generate a number of files entitled, for example “model_gas_pbtk”, but the only important one for *httk* is the .c file. Suggest copying the output file and adding “-raw” to the end of the unedited file name, before the .c extension to be consistent with naming conventions set in this paper. |
| 3 | Copy the .c file into the httk/src folder | This is the model file that will be compiled into the *httk* package when it is built. Remove “-raw” from the file name if you’ve already added it |
| 4 | Change the function names within the .c file | To be consistent with the naming convention, try to name the function, “[Function]_[Model Name]”. |
| 5 | Update the init.c file |  |
| 6 | Generate/modify .R files as needed |  |
| 6a | Generate/Modify modelinfo_[Model Name].R file |  |
| 6B | Modify parameterize_[Model Name].R file |  |
| 6C | Modify solve_[Model Name].R file | Depending on how many changes are made to the model itself, there may be little or no changes needed in the solve_[Model Name].R file |

To allow interoperability between the new model and the core HTTK functions in *httk,* a “modelinfo” file was created (Supplemental File “modelinfo_gas_pbtk.R”). Beyond basic definitions of the relevant functions, the modelinfo file describes the default units and state of matter (“gas” or “liquid”) for each compartment in the model. For the gas inhalation model these differed (for example, “exhaled breath” vs. “venous plasma”). Additionally, two new R functions were created for parameterizing the model and interfacing with the model solver (Supplemental Files “parameterize_gas_pbtk.R” and “solve_gas_pbtk.R”, respectively). In this case of gas_pbtk the code of the function parameterize_pbtk() was copied and modified, however it is also possible use parameterize_pbtk() itself to generate parameters for a variety of tissue lumping schemes. (For example, Kapraun, Sfeir (55) called parameterize_pbtk() twice within parameterize_fetal_pbtk() – once for the mother and once for the fetus.) The formatted C script (that is, “model_gas_pbtk.c”) was added to the “httk/src” sub-directory, and the three R scripts (that is, those files with the “.R” extension mentioned earlier in this section) were added to the “httk/R” sub-directory. Finally, the *httk* package was recompiled – as described in the ‘Methods’ section. This new PBTK model was included with the public release of *httk* (v2.0.0) [7].

Of the 1046 chemicals with *in vitro* measured data, only 962 are considered non-volatile or semi-volatile, and therefore within the applicability domain of the default “pbtk” model. The “gas_pbtk” model is applicable to all 1046. Linakis et al. added data for several dozen volatile chemicals to the internal httk data tables, and these data were also distributed with *httk* v2.0.0.

In Linakis et al.[7] a steady state solution was not developed. The lack of a steady state solution makes Monte Carlo simulation very expensive computationally and precludes many of the toxicological IVIVE tools built into *httk* (such as calc_mc_oral_equivalent()). However, the function calc_mc_tk() will run a series of full TK models (without steady state solutions) and report the mean and standard deviation of concentration at each time point requested.

### Model Evaluation

When new models are added to the suite of models in *httk* the incorporation of RMarkdown vignettes for reproducing key model evaluations provides a degree of quality assurance and invaluable documentation.

Linakis et al.[7] demonstrated an *httk-*based approach for evaluating generic TK models across multiple chemicals using *in vivo* measured data. The RMSLE was estimated for model predictions on 41 volatile organic chemicals across 118 experimental inhalation exposure conditions in two species drawn from the CvTdb [38]. Figure 1 was obtained by re-generating Figure 2 in Linakis et al.[7] by running the RMarkdown vignette “Linakis et al. (2020): High Throughput Inhalation Model” with the new features of *httk*. Figure 2 in that work compares observed chemical concentrations with predictions on a logarithmic scale. Three regressions were performed: separately for human and rat data, and then combined for an “overall” trend. In that study, prediction errors were analyzed across all chemicals jointly, so that chemicals with more data had greater weight. Since the available data can greatly vary between chemicals, we generally recommend first averaging per chemical, then averaging across chemicals, but given the aim of reproducing the previous work we do not do that here.

**Figure 1:**
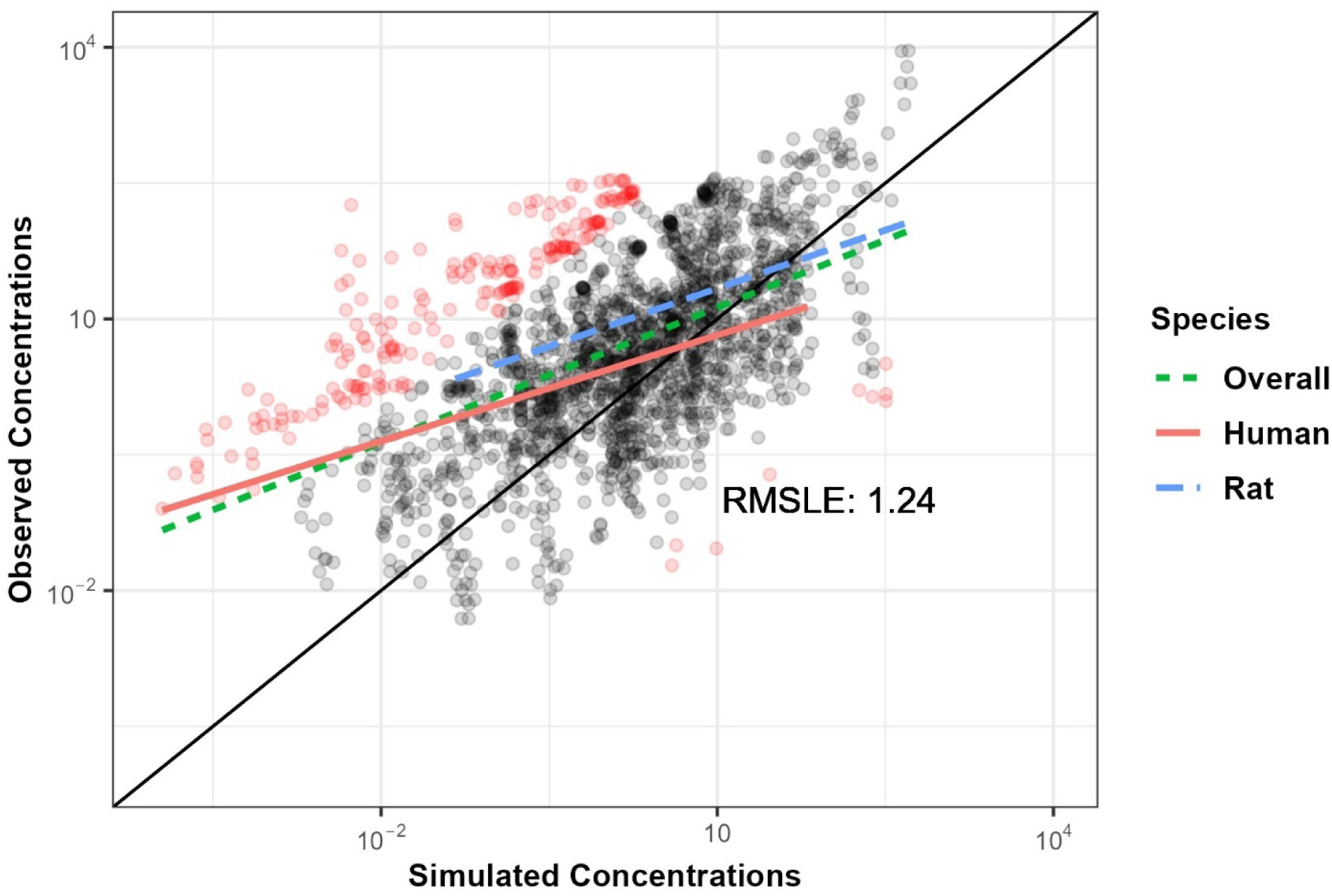
Updated evaluation of Linakis et al.[7] inhalation model using data from 118 in vivo experimental conditions in 2 species on 41 volatile organic chemicals obtained from the Concentration-vs-Time database (CvTdb) [38]. Black points indicate predictions within ten-fold of the observations, red points indicate more than hundred-fold difference. Line dashing indicates regressions between observations and predictions for humans (solid), rats (long dash), and both (short dash).

**Figure 2:**
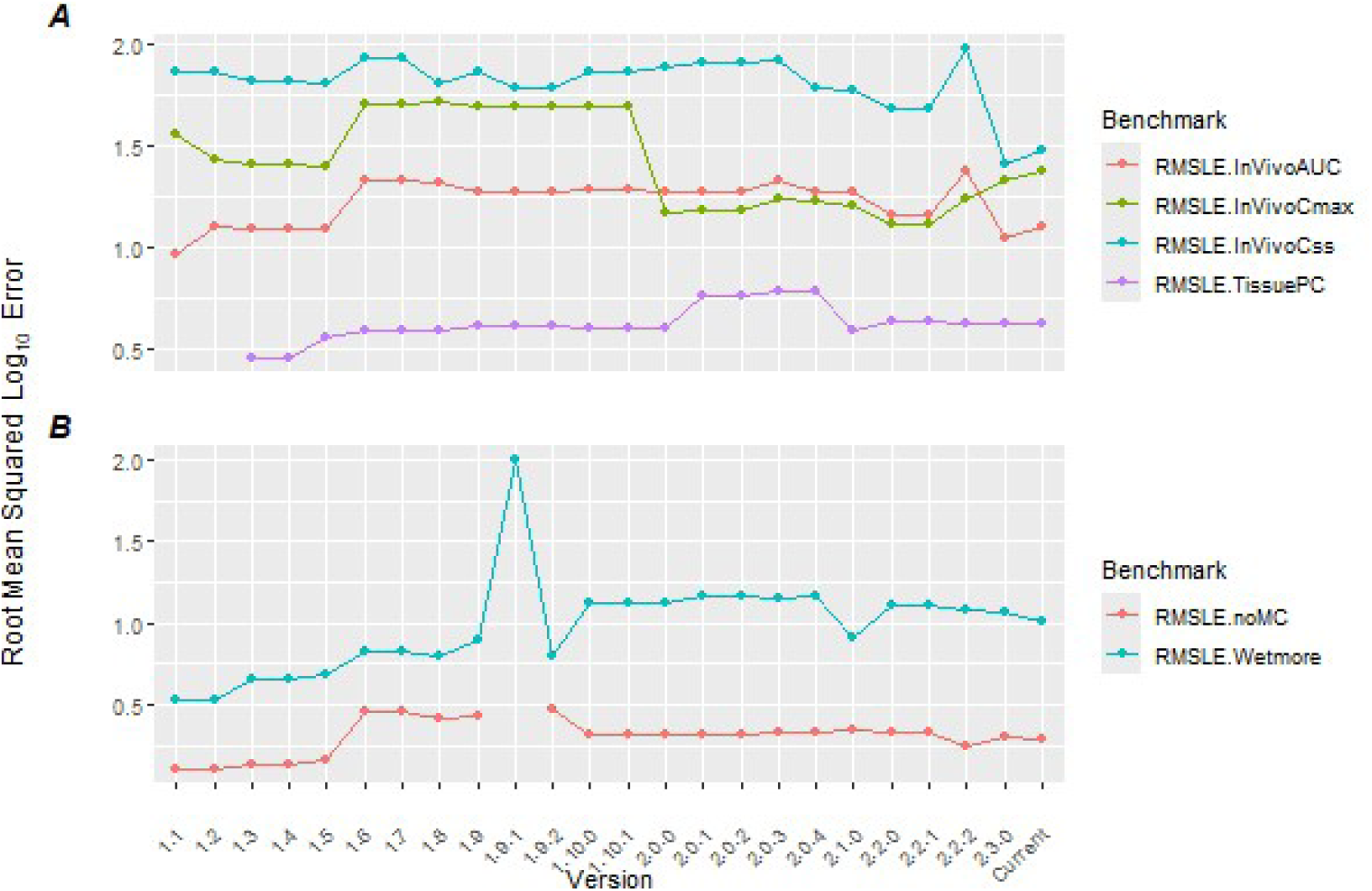
Panel A: Output generated by benchmark_httk() for both past and current performance of httk versions for a series of tests. Here performance is characterized by the root mean squared error on the log10 scale (RMSLE). IVIVE Statistics: In vivo area under the plasma concentration vs time curve (AUC), maximum observed plasma concentration (Cmax), and plasma concentration at steady state (Css) are all based on data collected by Wambaugh, Hughes (57). “Tissue PC” compares predicted tissue partition coefficients with literature in vivo estimates collected by Pearce, Setzer (58). Panel B: Wambaugh, Hughes (57). Monte Carlo Tests: The “noMC” test compares the median of the Monte Carlo C_ss_ obtained with calc_mc_css() distribution with the C_ss_ predictions of calc_analytic_css(). The “Wetmore” RMSLE compares the 95th percentile predictions of calc_mc_css() with the estimates from SimCyp [29] as reported by Wetmore, Wambaugh (27), Wetmore, Wambaugh (59).

Changes in results between our Figure 1 and Linakis et al.Figure 2 demonstrate the impact of incorporating new features within httk on the inhalation PBTK model. Here we observe that the RMSLE has increased from 1.11 to 1.24. This small discrepancy may be the result of several updates including: 1) restriction of model domain to non per- and polyfluorinated alkyl substances, 2) refinement of physico-chemical properties, and 3) on-going curation and refinement of the *in vivo* data within the CvTdb. R package “deSolve” [56] takes advantage of the speed of compiled C code (src/model_gas_pbtk.c) to calculate the derivative of the model ODEs to obtain rapid analytic solutions. The indices for the equations (indicating which variable is which) must match within both the C code and the R code. Within the modelinfo_gas_pbtk.R file the variable derivative.output.names lists the expected outputs, and must do so in the same order as the C code. For example, when developing the model we initially misordered the labels for variables that provide different units (for example, exhaled breath concentration is provided in both ppmv and uM). This mistake caused the RMSLE in Figure 1 to increase to nearly 1.8.

### Model Performance Benchmarks

EPA *httk* quality assurance procedures require that a new model be evaluated and accompanied by a peer-reviewed manuscript prior to formal inclusion into the CRAN distributable version of *httk*. Model evaluation is often the most difficult part, as it requires compilation of data amenable to perform the evaluation – for example, *in vivo* observations for chemicals relevant to the scenario explored by the model. Sometimes these data do not exist but, where they do exist, statistics indicative of a model’s predictive ability, such as RMSLE, absolute fold error, and the coefficient of determination (that is, R^2^), should be estimated. For R^2^ we suggest using the total explained variance (predicted vs. observed) rather than the R^2^ for a regression of the observations on predictions. There is no prescribed threshold of accuracy for inclusion to *httk*, though a case must be made that resulting model predictions are fit for purpose. Performance metrics for model predictability also serve as benchmarks for whenever functions within the R package are updatedand/or new data are added. We anticipate a general trend toward more accurate model predictions, though this may not always occur.

The new function benchmark_httk() performs a series of “sanity checks” that help assess the state of the package (Table 7). For example benchmark_httk() reports the number of chemicals with available data (which should not change unexpectedly. An additional sanity check is that outputs are generated for units of both mg/L and uM, and then the ratio 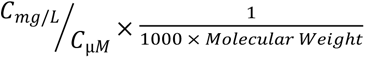 is calculated (this should be 1). benchmark_httk() also provides predictive performance benchmarks to assess the impact of changes to data, models, and feature implementations in the R package. In the past, some code changes made for one feature of *httk* have unintentionally impacted others. Most notably are the unit conversion errors introduced with hard-coded calculations (v1.9, v2.1.0). Package evaluation with this new benchmarking tool is meant to identify errors prior to version release thus reducing the impact on future users..

Performance of older httk versions was retroactively evaluated by manually installing previous versions of the package from https://cran.r-project.org/src/contrib/Archive/httk/, adding the code for benchmark_httk() at the command line interface and calculating the various benchmark statistics. Figure 2 shows the RMSLE as the benchmarking metric across several common HTTK statistics – including three IVIVE statistics, two Monte Carlo tests, and a partition coefficient test.

The *in vivo* statistics in benchmark_httk() establish the overall predictivity for area under the concentration vs time curve (AUC), peak plasma concentration (C_max_), and concentration at steady state (C_ss_) *in httk*. The *in vivo* statistics are currently based on comparisons to the *in vivo* data compiled by Wambaugh, Hughes (57). A summary of changes to *httk* by version are available at: https://cran.r-project.org/web/packages/httk/news/news.html

The partition coefficient test (“TissuePC”) provides an important check on the *httk* implementation of the Schmitt (60) model tissue to plasma equilibrium distribution. Pearce, Setzer (58) collected equilibrium tissue:plasma concentration ratios across a range of diverse chemicals and tissues. These predictions heavily rely on accurate descriptions of tissue composition and the ability to predict the ionization state of compounds being modeled. Therefore, the ability to predict tissue partitioning is particularly sensitive to changes in how physico-chemical properties are modeled by *httk*.

Monte Carlo benchmarks include the Wetmore and noMC tests. The “Wetmore” benchmark compares the calc_mc_css() Monte Carlo 95th percentile steady state plasma concentrations for a 1 mg/kg/day exposure against the steady state values calculated by SimCyp [29] and reported in Wetmore, Wambaugh (27), Wetmore, Wambaugh (59). These have gradually diverged as the assumptions for *httk* have gradually shifted to better describe non-pharmaceutical commercial chemicals [39]. The “noMC” benchmark compares the median of the distribution from calc_mc_css() against the non-varied steady state result from calc_analytic_css(). In both cases, deviation between the two values is expected but should not deviate wildly unless major changes have been made to *httk*’s Monte Carlo sampler or the underlying data. Notably in Figure 2, panel B, a unit conversion bug was introduced in the code for *httk* v1.9.1, altering several key results. The benchmark_httk() function was not available at the time, but with its implementation errors like this one can be identified and corrected prior to new version releases.

Both new models as well as other changes to the underlying *httk* code can generally be benchmarked against existing *in vivo* data using benchmark_httk(). Developers of new models may want to create and use *in vivo* data sets to benchmark new models against more relevant data (for example, characterizing new routes of exposure).

## Discussion

HTTK methodology predicts human doses with the potential to cause bioactive concentrations estimated from *in vitro* high throughput screening (that is, quantitative IVIVE [61]). One goal of the *httk* project is to add new TK models exploring various factors that may impact exposure. For example, new TK models might enable modeling for various exposure scenarios, physiological processes, or non-terrestrial species (such as fish) to better perform *in vitro* to *in vivo* extrapolation. The *httk* package is intended to maximize the reuse of existing *in vitro* measurements of TK determinants, physico-chemical property predictions, and functions to perform routine calculations with those values. By harmonizing functions and conserving data across HTTK models we aim to increase the verification of data and models. We also aim to improve package functionality, while improving transparency, reproducibility, and validity of new models.

The R package *httk* collects open-source data and models to both make HTTK available for chemical risk evaluation and make the HTTK methods accessible for review and refinement by the scientific community. The U.S. EPA’s CompTox Chemicals Dashboard [62], the U.S. National Toxicology Program (NTP) Interagency Center for the Evaluation of Alternative Toxicological Methods (NICEATM) Integrated Chemical Environment (ICE) [63], the European Food Safety Authority’s TK-Plate [64, 65], Health Canada’s “Bioactivity Exposure Ratio: Application in Priority Setting and Risk Assessment” science approach document [43], and Certara’s SimRFlow for SimCyp [66, 67], all make use of the existing data and models from *httk*. We hope that the data and simulation functionality made available by *httk* will encourage the modeling community to develop new HTTK models to expand IVIVE.

HTTK’s primary intention is for high-throughput human health risk prioritization. Models must be fully evaluated to be considered for use in public health for decision making [24, 68–71]. Clark, Setzer (25) identified six areas to be assessed for any PBTK model intended for public health risk assessment: 1) purpose, 2) structure and biology, 3) mathematical descriptions, 4) computer implementation, 5) parameter values and model fitness, and 6) any specialized applications. Ideally when new tissues or physiology are added to the *httk* modeling suite data characterizing these aspects will be used and made available to the HTTK community. In the absence of such data, new models should at least examine whether performance has degraded for other tissues (such as blood) where data are available.

R package *httk* facilitates systematic (statistical) evaluation using *in vivo* data for a range of chemicals. This evaluation allows quantification of uncertainty and characterization of confidence for decision making. The use of generic models in a standardized environment provides a variety of well-defined physiologies with specific routes of exposure that may be evaluated for many chemicals Thus, the first four points of Clark, Setzer (25) are addressed by *httk*. Issues of parameter appropriateness and model fitness require further evaluation using tools such as statistical regression and machine learning [48]. The R statistical programming environment provides many methods for this evaluation. Databases, like CvTdb [38], provide *in vivo* data to which model predictions may be compared for model evaluation. Often, the domain of applicability may be estimated empirically by correlating model performance and chemical properties [72].

Because the generic model describes a single physiology and characterizes chemicals with a set of standard descriptors, we only expect the model to perform well for chemicals where the included physiology and descriptors are relevant. If a process or property that is key to a chemical class is omitted then we expect the model to perform poorly for that class [37, 39]. While we expect larger uncertainty for any one chemical, a generic model allows greater confidence in model implementation [37] [30] and better understanding of trends across large numbers of compounds. We can consider the appropriateness of using a model based on *in vitro* data for chemicals without *in vivo* data if the magnitude of uncertainty and chemical properties are found to be statistically correlated [73].

The “software ecosystem” [74] design is intended to allow reuse [75] of data and functions (that is, recurrent calculations) from peer-reviewed scientific literature. As with any R package [52], reusable functions make *httk* more modular [76] and enables model developers to integrate new models and conduct consistent analyses. Developers may also extend existing functions to meet their needs. Function modularity provides “scoping,” which helps to limit the unintended impacts of new features and updates that developers incorporate in existing functionality [77]. Additionally, modularity centralizes code updates to the fewest places possible when errors occur, that is computational/code “bugs” in recurring calculations. Though these practices allow flexibility while attempting to prevent new contributions from breaking existing functionalities, bugs in the code will inevitably occur. It is critical for identifying and diagnosing issues that developers be able to conduct method and model evaluations that provide indications of a bug not identified by other means [70].

In Table 1 (and the Methods section) we describe how new models may be created and integrated with *httk*, and how previously evaluated tools and data may be used for new HTTK modeling approaches. Ultimately, the challenge is not building a new model, but rather evaluating the new model for its ability to provide accurate predictions under new conditions (for example, different exposure routes). New generic TK models can and should be statistically evaluated across the growing library of chemicals included in *httk*. Each new HTTK model added to the *httk* package should include an evaluation of model predictions relative to available *in vivo* data. Decision makers can then consider whether the generic model predictions based on *in vitro* data can be used to extrapolate internal doses for a chemical without *in vivo* data.

Though we can integrate new models into *httk*, there are still some limitations to the current tools provided in the R package. For example, there is currently no feature to check mass balance of the model prediction data output from solve_model(). Other future development directions include establishing methods for model operations (that is maintaining and evaluating HTTK models), addition of new datasets, streamlining access to data by functions within *httk*, improving the implementation of Monte Carlo simulation for greater flexibility when extending a new model to have uncertainty/variability propagation, and adding functions to continually standardize and streamline the process of adding new generic models to the *httk* R package.

The European Union Registration, Evaluation, Authorisation and Restriction of Chemicals has already begun the transition away from *in vivo* testing [78]. In 2018 the U.S. Environmental Protection Agency announced plans to phase out animal experimentation by 2035 [79]. *In vitro* new approach methodologies (NAMs) will be a cornerstone of next generation risk assessment [80]. Placing the results of NAMs in a human health context will require toxicological IVIVE for thousands of chemicals. This is the role for which *httk* is being developed [81].

## Methods

Reynolds and Acock (82) wrote that “modularity and genericness open models to contributions from many authors, facilitate the comparison of alternative hypotheses, and extend the life and utility of simulation models.” The same intention motivates the development of the *httk* R package for high throughput toxicokinetics. Table 1 provides step-by-step instructions or adding a new model into the httk environment, allowing that model access to all data distributed with httk, and hopefully allowing the model itself to be distributed to the scientific community. To ensure accurate description and evaluation of the models, *httk* developers use computer programming and modeling best practices such as version control software, unit tests, generalizable functions for creating a “software ecosystem”, and model evaluations. EPA uses the version control software Git [83] and maintains the codebase in a public GitHub repository (https://github.com/USEPA/CompTox-ExpoCast-httk) for transparent code tracking, including a history of changes over time and versions. Unit tests are part of general best practices when developing an R package [52, 84] (see “tests” sub-directory), and help determine if changes to one feature of *httk* unintentionally impact others by observing discrepancies between a set of expected and actual results. *httk* is freely distributed via the Comprehensive R Archive Network (CRAN – https://cran.r-project.org/) as a bundled “package” of R scripts, data files, and C code.

The remainder of this methods section describes in detail each element of how model developers can integrate their own generic HTTK models by using the modularity of *httk*. Additionally, we outline and describe some of the tools available to accomplish the addition of a new model including core *httk* functions, data, and other features (for example, model extensions). Though we mention specific functions, data, and features available in *httk*, this manuscript is not intended to provide an exhaustive list or description of *httk* functionality. Rather, specific tools mentioned are meant to demonstrate the utilization of key functionalities in *httk* for model integration or extension. For further details on functions, data, features, etc., we refer the reader to the ‘help’ files – which can be accessed via the help() function – vignettes, and other R resources to navigate the *httk* directory structure [52]. As with any R package, specific components of the package (for example functions and data) may be used without the rest of the package by adding the prefix “httk::” when developing applications or conducting analyses in R.

### Basic Generation and Integration of a New HTTK Model

This section reviews the basic parts required for incorporating a new HTTK model into the *httk* R package. Figure 3 provides a general schematic of the files and major components connecting the various software necessary for solving a typical HTTK model. Adding a new model to *httk* relies upon the use of three programs/coding languages – namely MCSim, C, and R. We will go through each part of the schematic as well as the fundamental files for integrating a new model into *httk*.

**Figure 3:**
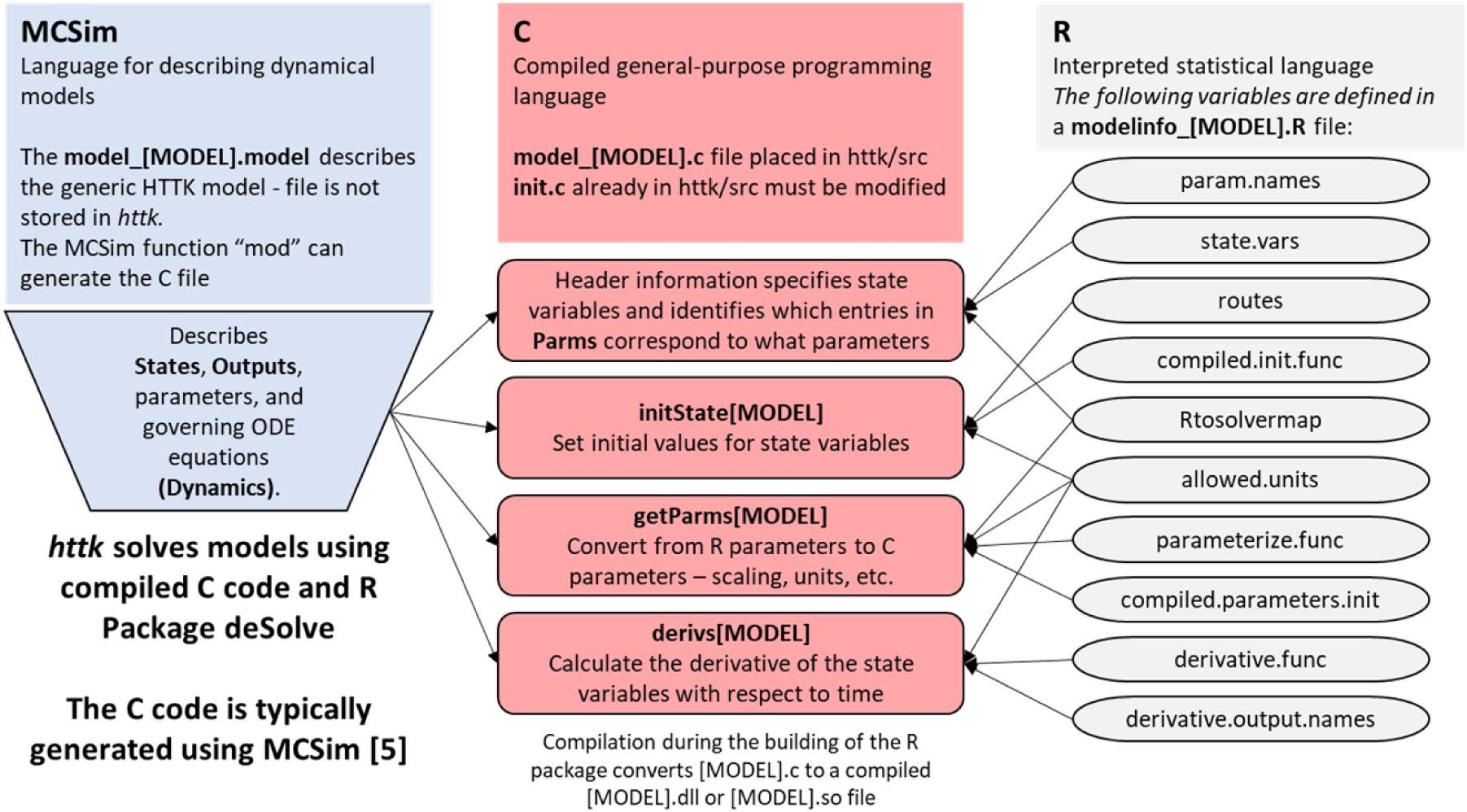
General schematic of the three software programs used to generate and implement a new generic HTTK model with the respective files, functions, and objects required for the three languages to communicate effectively. (Key files from left to right) The MCSim file, namely model_[MODEL].model, describing the ODE for the new HTTK model. The C file (updated), namely model_[MODEL].c, that deSolve will communicate with to evaluate the new HTTK model for a given exposure scenario. The model information R script, namely modelinfo_[MODEL].R, detailing information necessary for effectively preparing data to go into the HTTK model and handling results output from C via deSolve.

### HTTK Model File (MCSim and C)

The first necessary piece to incorporate a new generic HTTK model is the model description file. This file defines the set of ordinary differential equations (ODEs) for the time-evolution of the toxicokinetic model of interest. Here, we assumed the modeler uses MCSim [85], a dynamical modeling language, to define the HTTK model (model_[MODEL].model). MCSim was originally designed for all of the following: 1) implementing mathematical models (both ODE models and models that can be represented with algebraic equations); 2) performing MC simulations with mathematical models; 3) applying MCMC parameter inference methods to mathematical models; and 4) optimizing experimental designs. For *httk* we only take advantage of the first of these four applications of MCSim. Alternatively, modelers may write their HTTK model definition file directly in the compilable coding language C (model_[MODEL].c) and save it in the “httk/src” subdirectory of the *httk* package. In the latter scenario, this section may include some minor tips to consider when writing the model description file but can otherwise be skipped.

Generating the model definition file with MCSim (with extension “.model”) first requires the system of equations for the toxicokinetic model. *httk* uses MCSim to describe some, but not all aspects of the simulation. In particular, dosing (including initial conditions and later events) are handled within R. Table 2 provides the list of key components for defining an ODE model – necessary for appropriately defining and incorporating the model – and their mapping between MCSim (.model file), C code (.c file), the *httk* R package (modelinfo_[MODEL].R file), and the *deSolve* package (ode function call in function solve_model()) [56, 86] .

**Table 2:** Major components for describing an ODE model and their mapping between the modeling concepts used by MCSim, deSolve, and httk.

| ODE Components | Description | MCSim | C Code (httk/src) | R Code (httk/R) | deSolve |
| --- | --- | --- | --- | --- | --- |
| System of Differential Equations | The ODE system describes the evolution with time of the state variables as a function of the current state and model parameters. | MCSim includes a specific “Dynamics” block. These equations describe the derivative of the model ODEs. MCSim is an interpreted language so that the order of does not matter. | MCSim function “mod” translates the ODEs to C code for calculating the derivative. For C the values evolve as each line of code is executed. | modelinfo_[MODEL].R points to a function “derivative.func” that calculates the derivative of the ODEs. | For htkt, function deSolve::ode argument “func” is set to the name of a compiled function that returns a vector of derivatives for the system of ODEs describing the model. Derivative function is specified by variable “derivative.func” in the modelinfo_[MODEL].R file. |
| States | The state variables are those described by the system of ordinary differential equations (ODEs) | MCSim includes a specific “States” block. States are “variables for which a first-order differential equation is defined in the Dynamics section.” [87] | C arrays “y” and “ydot” contain the state variables and the derivative with respect to time | modelinfo_[MODEL].R describes “state.vars” which should be in the order of derivatives returned from “derivative.func”. | The first argument to function deSolve::ode, both defines initial values for state variables and implicitly the number of state variables. |
| Model Parameters (Non-state variables) | Each equation describing the time-evolution of a state variable may contain multiple parameters describing both physiological and chemical properties | List of parameters, named and assigned values where necessary in the MCSim code. That is, there is no explicit “Parameters” block, but rather they are collected by MCSim into a list based on wherever they show up. | C array “parms” contains all N parameters used by the model, the first value is parms[0] and the last is parms[N-1] | In the modelinfo_[MODEL].R file the variable “param.names” lists all the parameters used by R, which sometimes includes things such as logP, which underly the parameters used by the ODE system but are not directly needed by the model. “Rtosolvermap” identifies which R parameters link to C parameters described by “compiled.param.names”. As with States, these must be defined in the order used by C. | deSolve::ode takes an argument “parms” containing just the parameters needed by the derivative function “func”. |
| Times | Time points for simulation, including timing of events (such as dose administration) | Simulation times do not need to be explicitly described for our use of MCSim. | The derivative function is written to calculate the instantaneous derivative at time “pdTime” | For calls to htkt the “times” vector only needs to include the observed (measured) times – dosing times are | deSolve::ode argument “times” takes a vector of times where predictions are needed, though |
|  | and necessary prediction (such as measurements) |  |  | added automatically. Inclusion of intermediate time points can in some cases help the ODE solver by forcing it to take smaller steps. | intermediated times are calculated. The first value must be the initial time. |
| Initial Conditions and Events | Instantaneous dosing changes in state, such as an additional exposure to a chemical. These are handled by breaking the simulation into the time before and after the change. | We do not make explicit use of MCSim's Initialize or Inputs block, and instead use the functionality of deSolve. Initial values of state variables do not need to be explicitly described for our use of MCSim. | Events are handled outside the C code by *deSolve*. The C code describes the model between events. | R data.frame "eventdata" (determined from dosing.matrix, doses.per.day, and daily.dose if available) is constructed by `httk::solve_model` and passed to `deSolve::ode` as argument "event". | Events impose discontinuities in state variables at specific times. *deSolve* uses numerical integration algorithms to describe continuous changes in between events. At each event one or more state variables are changed instantaneously. [87] The first argument to function `deSolve::ode`, R vector "y", defines initial values for state variables. |
| Inputs / Forcings | Changes in other external factors (for example, chemical concentration in air or constant infusion dosing). May be used as an additional dosing parameter for particular exposure scenarios. | MCSim includes a specific "Inputs" block. Inputs are independent variables for the model ODE system. They may change over time due to Events or Forcings. [87] | C array "forc" contains the values of the forcing variables. | Determined from `initforc`, `forcings`, and `fcontrol` if available | *deSolve* treats the forcing function as a time series of constraints – the relevant state variables must be such that the value of the forcing function is always matched. [87] |
| Outputs | The ODE model components to export from the model evaluation. Changes in state variables must be returned. However, it may also include related variables such as concentrations (when state variables are amounts), state variables in different units, mass-balance checks, and time-integrated values (for example, area under the curve - AUC). | MCSim includes a specific "Outputs" block. They can be both state variables and functions of those variables. | C array "yout" may contain more than just the derivatives of the state variables, providing a means of returning values calculated from the state variables to *deSolve* or *httk*. | R data.frame `out` contains a row for each time and a column for time plus all variables requested via the "monitor.vars" argument. | *deSolve* function `ode` returns a data.frame with a row for each time requested and a column for each output returned by the C function |

For further details on the MCSim modeling language and more complex modeling needs, outside of what is described here and necessary for *httk*, we refer the reader to the MCSim documentation (https://www.gnu.org/software/mcsim/mcsim.html) [88].

Once we have the MCSim model description file we need to translate the “.model” file to a compilable C script, because R is unable to directly interpret MCSim files. C is a fast machine-specific language[89], which performs rapid model simulation and there is an existing package to allow for direct communication between R and C for dynamic modeling (deSolve [56]). Generating the compilable C script, which will be saved in the ‘httk/src’ sub-directory, from the MCSim file requires using the “mod” function from MCSim [88] and some minor updates to the initial C script. The following command executed in a command-lineterminal converts an MCSim file to a C file:

mod -R <NAME>.model <NAME>-raw.c

The generated C version of the model definition file (with extension “.c”) will also be accompanied by an initialization file (extension “_inits.R”). Note, the suffix “-raw” in the C file name (model_[MODEL]-raw.c), which denotes this file is an unformatted version of the model definition file. The “raw” C file will require some minor updates prior to being saved as the final C file (model_[MODEL].c) in the ‘httk/src’ sub-directory. Additionally, the “_inits.R” file(s) generated during the conversion may be deleted because they are unnecessary for the R package and the functionality is already provided elsewhere in *httk*.

As mentioned above, the C file model definition file requires a few minor modifications prior to integration into *httk*. The modifications to the function names created by MCSim help to avoid confusion amongst the suite of existing models in the package. Table 3 details the parameters and functions that require consideration, in which files they are found, and what updates are necessary.

**Table 3:**
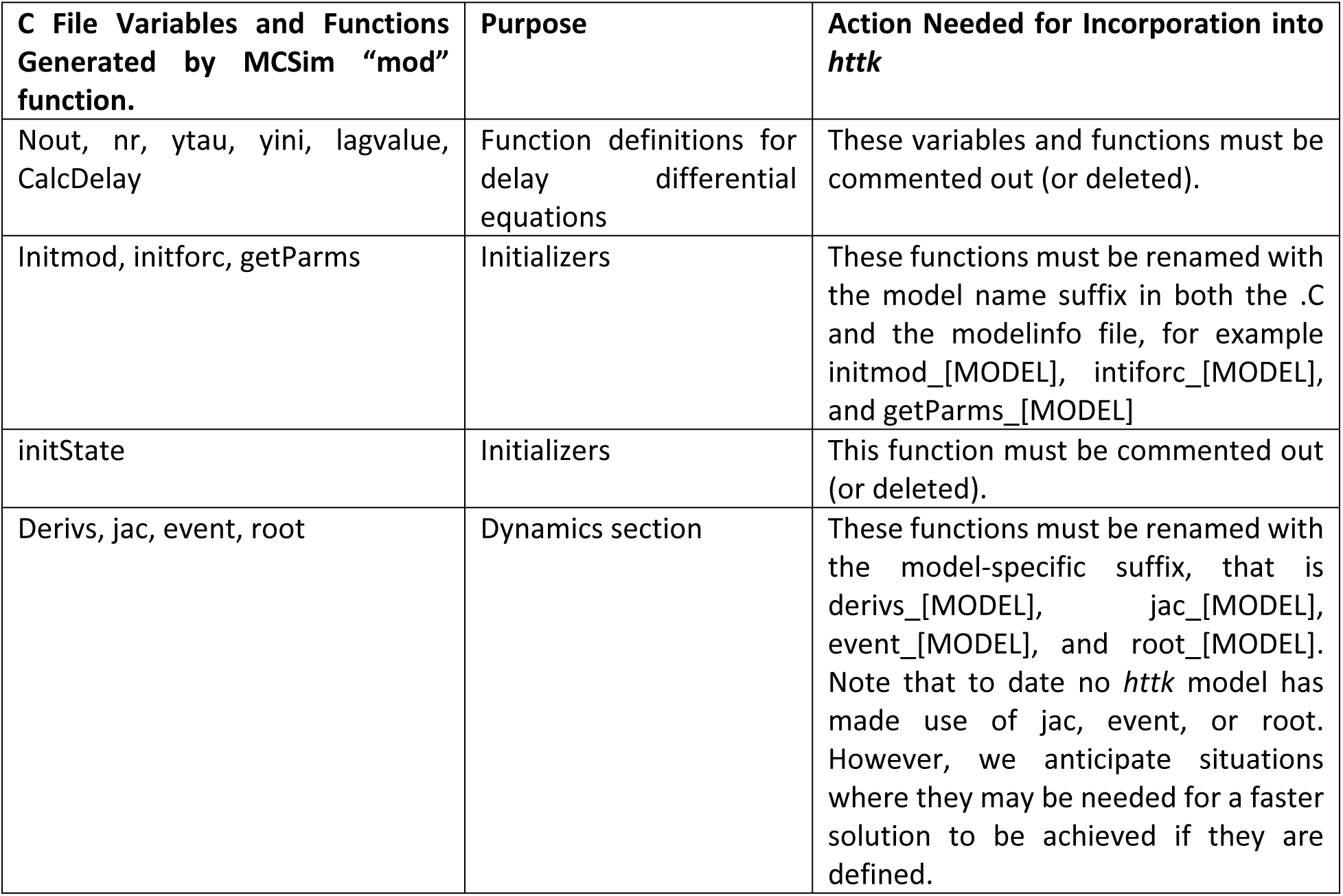
List of functions and variables auto-generated by MCSim’s “mod” function, when converting the model definition file from the MCSim language to C, that must be renamed or removed prior to incorporation into the R package “httk”.

First, we need to update the standardized function names, generated by MCSim to be model-specific. MCSim assumes the individual models are standalone and does not consider that they may become part of a suite of models located in the same directory. Model-specific names for functions and parameters are created by appending the model’s name as a suffix in the newly created “raw” C file. For example, the 3-compartment steady state model, abbreviated to “3compss”. The “3compss” is the model name suffix that will replace the generally referenced “[MODEL]” suffix found in the function names in Table 3. This designates that the functions and parameters are specific to the 3-compartment steady state model. The second modification to the “raw” C file requires removal, or commenting out, of unnecessary functions included as part of the MCSim to C code conversion. These include functions or parameters such as “Nout” or “initState”, see Table 3 for a full list. Once the necessary modifications are made to the C file we save a copy of it, without the “-raw” filename tag, to the ‘httk/src’ sub-directory.

### Model Specification & Parameterization Function Files (R)

After the C model definition file is created and saved in the R package, we then generate a model information file and any necessary parameterization functions (R scripts) based on the defined HTTK model. These R scripts will be stored in the ‘httk/R’ sub-directory of the *httk* package. Each of the R scripts described in this section allow model developers to incorporate their model with the basic functionality of *httk* as well as the more complex extensions (for example, steady state modeling or Monte Carlo simulations), while maintaining a standard approach using a set of generalizable ‘core’ functions provided in *httk*. Figure 4 provides a basic visualization showing the connection between the key auxiliary files – for example the model information, parameterization function, and core functions – enabling the incorporation of the new model into *httk*.

**Figure 4:**
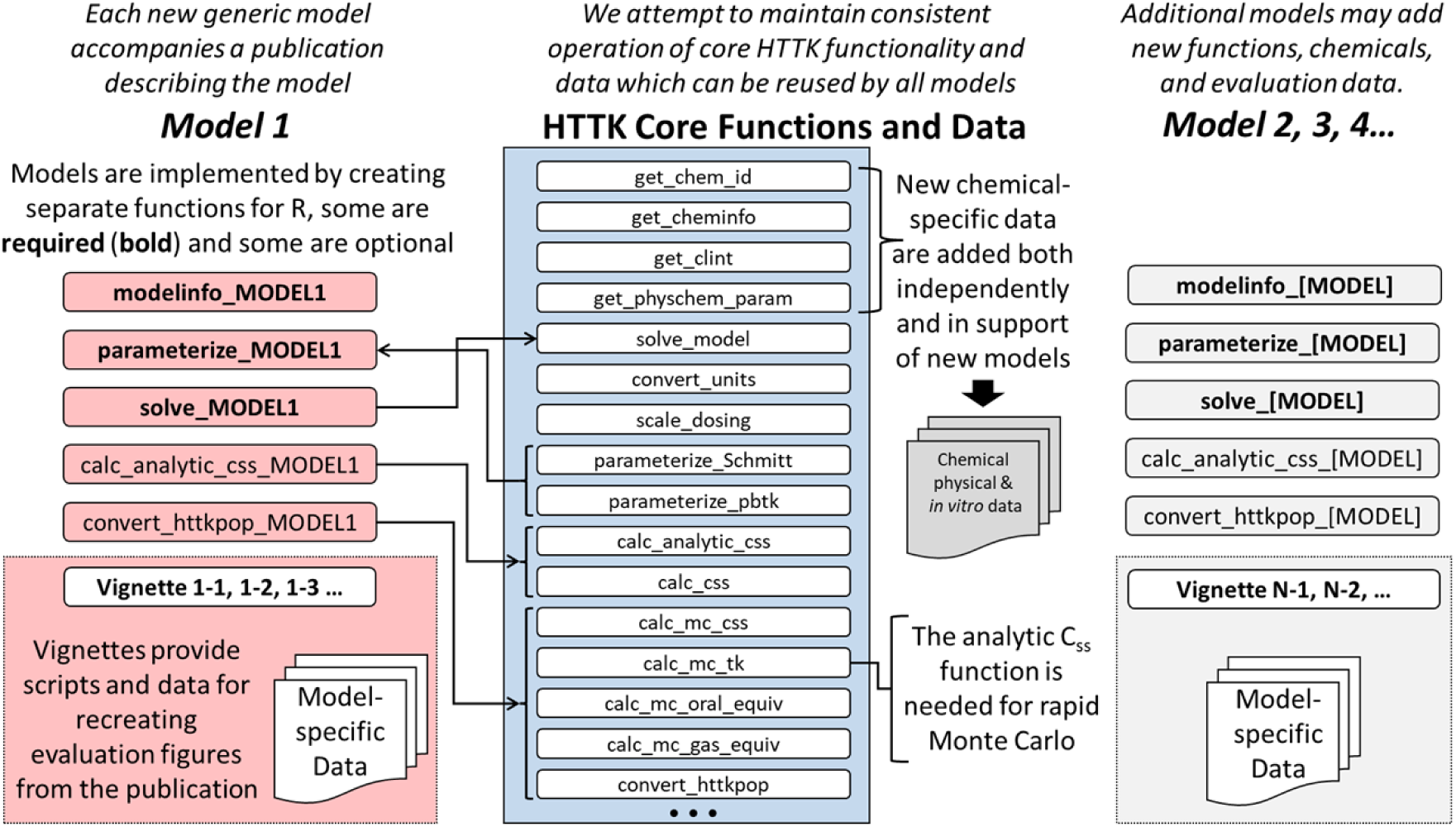
httk is intended to permit flexibility with respect to the description of toxicokinetics but allows the reuse of data and functions for tasks such as parameterization, unit conversion, steady state calculation, and Monte Carlo simulation. HTTK Core functions are defined in Table 5. Other items are defined throughout the text. We refer the reader to the Supplementary File S2_function_guide_file.xlsx for a full list of existing httk functions with brief descriptions and the help files within the httk R package for further details.

Version 2.0.0 of the *httk* package implemented a set ‘core’ functions providing a common interface to standard functions such as data reading, parameter entry/modification, and model execution, to generalize interactions with all the models hosted in *httk*. To facilitate these interactions, the model information file (modelinfo_[MODEL].R), more generally referred to as the “modelinfo” file, was developed. The modelinfo file acts as a reference guide for each HTTK model providing model-specific global variables, mappings between R and C variables and functions, as well as criteria indicating data that are relevant for model evaluation. Each modelinfo file saves the standard set of descriptors to the ‘model.list’ object within *httk*, which holds the model-specific information for all models in the package. The top level of list elements in ‘model.list’ correspond to a single generic HTTK model allowing it to act as a look-up object when ‘core’ functions need model-specific information to perform a task. The ‘model.list’ object allows the ‘core’ functions to identify and interact with all models in a standardized manner using standardized descriptors.

For a new generic HTTK model, the corresponding modelinfo file needs to create a new list element in the ‘model.list’ object, namely “model.list[[MODEL]]”. “MODEL”, as previously described, designates a generic placeholder for model-specific abbreviations. Basic model incorporation requires 1) defining the components that allow communication between the R and C languages, 2) tissue information, 3) applicable data information, and 4) and consideration of units. All the basic components are listed in Table 4.

**Table 4:** Basic ’model.list’ elements to include in the modelinfo file when incorporating a new generic HTTK model into “httk”. Note, this table only covers the basic model and data criteria components not any model extension components. We refer readers to the Supplemental File S1_modelinfo_guide_file.xlsx for further details on all components.

| Component Type | Model Specific List Components | List Component Functionality |
| --- | --- | --- |
| Basic/Dynamic | parameterize.func, solve.func | R Functions |
|  | alltissues, tissuelist | Tissue Information |
|  | param.names, required.params | R Parameters |
|  | Rtosolvermap | R to C Parameter Mapping |
|  | compiled.param.names, derivative.output.names | C Parameters |
|  | compiled.parameters.init, compiled.init.func, derivative.func | C Functions |
|  | allowed.units.input, allowed.units.output, compartment.units | Default Unit Information |
|  | default.monitor.vars, state.vars | C Variables |
|  | routes | Dosing Components |
|  | forcings.materials, input.var.names | Forcing Dose Components |
|  | compartment.state | HTTK Rout and Chemical State Components |
| Data | exclude.fup.zero, log.henry.threshold, chem.class.filt | Data Inclusion/Exclusion Criteria Components |

For model developers intending to extend their model with additional *httk* functionalities (for example, steady state calculations, uncertainty propagation, etc.) additional components will need to be included in the modelinfo file. However, we will leave this to be discussed in the Generic TK Model Extensions Section where other extension requirements are described. For additional details demonstrating how to create the modelinfo file we refer the reader to the *httk* vignettes and help files – see Supplemental File S1_modelinfo_guide_file.xlsx for list of possible components.

Once the modelinfo file is generated, the next focus is to create the parameterization function script (parameterize_[MODEL].R). The function in this script interacts with a larger set of functions and data in *httk* defining a series of steps to obtain and format *in vitro*, physiology, and physical-chemical property (physico-chemical) data necessary for parameterizing and solving the new HTTK model. In some cases, there is no dynamic modeling component (that is, no C script) and the parameterization function is the only necessary component to run the model (for example, parameterize_schmitt()). Regardless of the type of parameterization function being developed the essential information typically includes – but is not limited to – relevant physiology data, chemical identifiers, relevant tissues, and physico-chemical parameters.

While some models use their own approach for retrieving chemical-specific model parameters and generating model parameterization data, many are developed around an existing core function parameterize_pbtk(). The parameterize_pbtk() function can be customized to create a model-specific function by specifying which tissues are lumped together into aggregate compartments. For example, parameterize_3comp() makes use of parameterize_pbtk() to obtain all necessary parameters and sets any unnecessary parameters to NULL. More complex models may use parameterize_pbtk() several times in their model-specific parameterization function. One such model is the maternal-fetal model from Kapraun et al. (2022) [55], with the corresponding parameterize_fetal_pbtk() function that calls the parameterize_pbtk() function twice. The first call obtains parameters for the mother and the second call obtains parameters for the fetus. However, if a modeler needs a more flexible approach from calling existing parametrization functions, they may consider calling other core functions that accomplish similar tasks – for example get_chem_id(), get_clint(), get_physchem_param(). A modeler may need to include core functions and data as part of the parameterization function which are discussed in more detail in the General Utility Core Function Section. We refer readers to the *httk* vignettes for further details on creating a parameterization function for the new HTTK model, which may have a unique set of needs. The modelinfo and parameterize function R scripts should be saved in the ‘httk/R’ sub-directory of the *httk* R package once they are created.

### Wrapper Function for the New HTTK Model

In addition to the modelinfo and parameterization function R scripts, developers may want to automate running their new generic model. Most HTTK models have a “wrapper” function (solve_[MODEL].R) calling the generic solve_model() function and tailoring arguments for the model, for example solve_1comp(). “Wrapper” functions generally configure, evaluate, and/or extend basic functionalities of the primary analysis by encapsulating it with recurrent steps used before or after the primary analysis/function call – for example, data retrieval and formatting. Some recurrent steps can be accomplished using core functions, such as pulling physico-chemical data or performing unit conversions. Generating a wrapper function allows the model prediction process to be generalizable to the extent possible given model constraints and applicability. It enables evaluation across a variety of scenarios, standardization of the process, and potential for mitigating errors. Though creating a wrapper function is not required for specific HTTK models, it does have two major advantages. First, it allows for more reproducible results, depending on the function configuration. Second, it encourages more wide-spread application of the new model by making it readily available for *httk* users that are not necessarily modelers but want to make use of the model’s predictions for their own analysis.

As mentioned previously, solve_model() is a generic function used to communicate with the compilable C form of the HTTK models through R package *deSolve* [87]. *deSolve* interfaces between R and C to obtain a solution to the system of ODEs. solve_model() is meant to be generic and usable for any HTTK model. Therefore solve_model() is designed to accept systematized data and metadata for a given toxicokinetic model. This includes – but is not limited to – variable names, parameterization functions, and key units. When this model description information is passed to solve_model(), along with chemical-specific descriptors, the function can set up the ODE model system to obtain the numerical solution of chemical amounts (or concentrations) in different body compartments over time. Much of the information solve_model() requires is either specified in the modelinfo file – described previously – or pulled from *httk-*provided datasets using core functions like get_cheminfo() or get_physchem().

It is imperative that the data provided to solve_model() and passed to the C model are in the units anticipated by the model. Thus, the convert_units() and scale_dosing() functions are commonly employed within wrapper functions. This allows users to provide their experimental data “as-is”, that is without any unit conversion, to the wrapper function and the unit conversion and scaling functions ensure the data are appropriately converted to the expected units in a seamless and repeatable manner. For further details on constructing a wrapper function, namely solve_[MODEL](), we refer users to the *httk* vignette demonstrating how to create and document theat function along with some considerations when developing a new HTTK model.

Once the necessary files are generated and saved in their respective sub-directories we need to compile and install the package (“R CMD INSTALL httk” in the operating system command shell) prior to execution. Then we can obtain concentration vs. time predictions for a given set of chemicals with our newly incorporated model. Re-installing the package with a new model allows use of existing functionalities and package dependencies to solve the ODEs in the new generic HTTK model. Once compilation and installation are completed, we can run our newly incorporated model to obtain concentration vs. time predictions for a set of chemicals.

### General Utility Core Functions

Throughout the past few sections, we gave a high-level description of the process for compiling and incorporating a new generic HTTK model into the *httk* R package. In this section, we highlight some of the core functions in *httk* that enable modularity and harmonization of models across the package. Core functions are generalized, yet ‘model-aware’, and can be used with any generic HTTK model added to *httk* (see Figure 4). The model-aware attributes of core functions allow for tailored results based on the model with which they are interacting.

Consider, for example, that each generic HTTK model requires different parameters. Therefore the chemicals with sufficient descriptive parameters may vary model-to-model. In this instance, the get_cheminfo() function serves to identify and report chemicals applicable for the specified model. For example, if your model requires fraction unbound and intrinsic clearance, get_cheminfo() will only identify the chemicals that have those parameters available. Optional arguments in the core functions give users the flexibility to tailor results for their needs. Continuing with our example function, get_cheminfo(), the “info” argument defaults to returning only the CAS (Chemical Abstracts Service) identifiers, but a user may also specify other data to include. To include additional information, users provide a character vector specifying additional data to retrieve – for example, logP and molecular weight. Another optional argument necessary for some models is “class.exclude”, which checks the model information file to see if there are any chemical classes that are outside the domain of applicability for the specified model and ought to be excluded. Though we will not provide an exhaustive list here of all available core functions in *httk,* and their arguments, Table 5 outlines several commonly used functions with a brief description of their utility. For additional details we point the reader to the *httk* help files and Supplemental File “S2_function_guide_file.xlsx” which have an exhaustive list of available functions.

**Table 5:** Reusable Core Functions with General Utility for Generic TK Models. (Note, this table only lists a subset of core functions available in the httk R package and is limited to functions explicitly mentioned in this paper. We refer readers to the Supplemental File S2_function_guide_file.xlsx and help files for further details on these and other available functions.)

| HTTK Core Function | Description |
| --- | --- |
| `get_chem_id()` | Interface for users to check if a chemical of interest is available within *httk* data and returns all three possible chemical identifiers (that is, chemical name, CASRN, and DTXSID). |
| `get_clint()` | Interface for users to retrieve chemical- and species-specific intrinsic hepatic clearance ($Cl_{int}$) for a given list of chemicals. |
| `get_physchem_param()` | Interface for users to retrieve physico-chemical properties from the `chem.physical_and_invitro.data` table in *httk* for a given list of chemicals. |
| `get_invitroPK_param()` | Interface for users to retrieve *in vitro* pharmacokinetic (PK) data, for example intrinsic metabolism clearance or fraction unbound in plasma for a given list of chemicals and species. |
| `get_cheminfo()` | Interface for users to list the chemicals with sufficient data to run a specified HTTK model. Interacts with internal HTTK data tables containing chemical-specific information. [NOTE: By default, the function assumes the model is “3compartmentss” and reports the chemical identifiers as Chemical Abstract Services Register Number (CASRN).] |
| `add_chemtable()` | Utility function that adds chemical-specific information to the `chem.physical_and_invitro.data` table necessary to parameterize a model, such that additional chemicals can be modeled/evaluated. |
| `solve_model()` | Generic numerical ODE solving function that uses *deSolve* to call the C code for the derivative and simulate a concentration vs. time response. Often used by wrapper functions (for example, `solve_steadystate()` and `solve_pbtk()`). |
| `convert_units()` | Utility function for converting input and output values from their current units to the appropriate units necessary for the model or user interpretation, respectively. |
| `scale_dosing()` | Utility function converting dose in terms of unit per body weight to specified dose units not in terms of body weight. |
| `parameterize_schmitt()` | Utility function providing the necessary parameters to estimate predictions for plasma partition coefficient [58, 60, 90]. |
| `parameterize_pbtk()` | Utility function generating a set of chemical- and species-specific PBTK model parameters, including tissue:plasma partition coefficients, organ volumes, and flows for lumping tissues. Tissues should be described in the `tissue.data` datatable. |

The two most crucial core functions mentioned in Table 5, which users should use when incorporating a new model, are the convert_units() and scale_dosing() functions. These functions are crucial because they implement tools that adhere to the “Rules of PBTK Modeling” (Dr.s Mel Andersen and Harvey Clewell, among many who started out at the Wright-Patterson Air Force Base in the United States, have spent decades realizing the potential of PBTK to inform chemical safety both through research and educational outreach. Their classes often feature these rules):

1. Check Your Units
2. Check Your Units
3. Check Mass Balance

Unit conversions are among the most common sources of PBTK model errors [25]. Thus, the convert_units() and scale_dosing() functions aim to mitigate this source of error by defining the relevant calculations in one function, and calling the relevant function when a conversion is necessary. The scale_dosing() function performs species-specific body weight scaling by detecting “/kg” in unit strings, which indicates units in per kilogram body weight, then scales the dose values with the respective multiplier (body weight). Alternatively, the convert_units() function provides conversions for many other units, specifically amounts and concentrations (that is, amount per unit volume). For example, convert_units() can be used to convert “mg” to “µmol”, or “mg/L” to “µM”. It should be noted, for conversions between concentrations and amounts – for example “mg” to mg/L” – a volume value is required. Implementing reusable unit conversion functions minimizes the burden on users to manually include these calculations in their code and avoids the unintentional introduction of bugs due to typographical errors. More importantly, it centralizes the code where these conversions are defined. This greatly facilitates evaluation of the units used in inputs, in calculations and in producing results such as figures. We refer readers to *httk* help files and vignettes for further details on using these functions, how to add new conversions, and other details to consider.

### Data in *httk*

In addition to the core functions, *httk* also provides a set of data tables with information needed to parameterize HTTK models. These tables help enable modularity and consistency by eliminating the need to hard code model parameter values. There are several key datasets used to parameterize and evaluate generic HTTK models. These datasets are stored in the Tables.RData file under the “httk/data” sub-directory.

Table 6 provides a list of core datasets that are automatically loaded into a local R session when *httk* is called via “library(httk)”. These include, but are not limited to, two chemical and two physiological parameter tables as well as a metadata object with information about the date of the data table generation. Users can retrieve parameter values stored in these tables with the corresponding core functions. Though there are other tables available in the package, aside from those listed in Table 6, they are not considered part of the core set necessary for parameterizing and solving HTTK models. Here, we describe the four core datasets and their role in incorporating a new model in *httk*. We refer the reader to the *httk* vignettes and help files for details about using these and all other available datasets.

**Table 6:** Core HTTK Data Sets – includes chemical specific tables with physico-chemical properties, physiological data, and data generation meta-data.

| Data Object | Description |
| --- | --- |
| `chem.physical_and_invitro.data` | Chemical-specific values for physico-chemical properties and *in vitro* measurements. |
| `chem.invivo.PK.data` | Chemical concentration vs. time database (CvTdb) snapshot for evaluating model predictions [38]. |
| `physiology.data` | Data describing species-specific physiology. |
| `tissue.data` | Tissue composition data for Schmitt’s method of partition coefficient prediction. |
| `Tables.Rdata.stamp` | Time stamp identifying when the other tables were created. |

### Chemical Specific Data

The chemical-specific *in vivo* and *in vitro* measurements as well as the physico-chemical properties in dataset chemical.physical_and_invitro.data make it the most used parameter table. chemical.physical_and_invitro.data contains many of the parameters vital for estimating absorption, distribution, metabolism, and elimination from the body after an exposure – for example molecular weight or hydrophobicity (that is, logP). There are four key types of chemical-specific parameters available for use with HTTK models – including hepatic clearance (Cl_int_), fraction unbound in plasma (f_up_), blood-to-plasma ratio (Rblood2plasma), and membrane permeability (oral absorption as characterized by Caco-2). More detail on these parameters is provided at the end of this section.

Though all the *httk* functions retrieve chemical-specific data from chem.physical_and_invitro.data, few of them directly interact with that data table. There are a cascade of decisions for selecting the appropriate value from chem.physical_and_invitro.data. Decisions include identifying the appropriate species-specific value (or surrogate) and whether *in vivo* or *in vitro*-derived values are to be used. Several functions provide an intermediate layer interacting with chem.physical_and_invitro.data, including: get_cheminfo(), get_invitroPK_param(), get_physchem_param(), and add_chemtable(). The first three functions are useful when a new model needs to pull chemical-specific data as it does this consistently. The fourth function, add_chemtable(), allows a modeler to insert additional data that may be needed to evaluate the new model. add_chemtable() is for adding data about new chemicals – adding new descriptors (such as physico-chemical properties) currently requires modifying the add_chemtable() function itself.ki We refer the reader to the package help files for further details about using these functions when developing their new model.

Most of the *in vitro* data are measured from human tissues. However, for certain calculations a user may want to use values from non-human physiologies –for example, rat, mouse, etc. – to solve a model. The “default.to.human” argument is provided for many of the parameterization functions to allow this option. In cases where the human values are preferred in lieu of non-human species, users should set “default.to.human” to TRUE.

Intrinsic hepatic clearance (Cl_int_), for the purposes of *httk*, is an *in vitro* measurement for the rate of disappearance of a compound when it is incubated with hepatocytes [91]. Cl_int_ is interpreted as a measure of metabolic rate by the liver, which for many chemicals is the primary organ of metabolism. In *httk,* Cl_int_ is stored in units of µL/min/10^6^ hepatocytes. When scaled appropriately, Cl_int_ is predictive for the clearance rate of a compound from the body via the liver. Cl_int_ adjustments are executed via the calc_help_fu() function in *httk*, which accounts for free fraction of chemical in the hepatocyte assay [92]. The calc_hep_clearance() function uses Cl_int_ values to estimate whole-liver hepatic clearance based on various scaling models described by Ito and Houston (93), typically assuming the well-stirred model when the liver is not modeled as a separate compartment. The Cl_int_ parameter may be reported as a single value or a 4-tuple string of values separated by commas. For 4-tuple string of values, these are Bayesian estimates where the median is first, followed by the lower and upper 95^th^ percentile values, and concluded with the p-Value for systematic clearance (that is, likelihood that there was no clearance observed), respectively. If users extends their model with the Monte Carlo sampler, discussed in the Monte Carlo Simulation section, this information will be used, when available, to appropriately propagate chemical-specific uncertainty throughout the model.

Fraction of chemical unbound in plasma (f_up_) is a central parameter for estimating other predictors within a generic HTTK model. There are data from two different *in vitro* assays measuring f_up_ included in *httk*: rapid equilibrium dialysis [94] and ultracentrifugation [95]. *httk* assumes f_up_ is constant across time and throughout the body. The f_up_ parameter may be reported as a single value or a 3-tuple string of values separated by commas. For 3-tuple string of values, these are Bayesian estimates of f_up_ where the median is first, followed by the lower and upper 95^th^ percentile values, respectively. Finally, f_up_ may require adjustments to account for differences between available lipid amounts *in vitro* vs *in vivo* [58]. These adjustments can be made using the calc_fup_correction() function.

Oral absorption is the last chemical-specific parameter necessary for many of the *httk* models. This parameter is used for estimating the amount of a chemical taken up in the body via ingestion (oral exposure). Honda et al. (submitted) describes *in vitro* measured Caco-2 permeability data and a quantitative structure-property relationship model for estimating chemical absorption. The function get_fabsgut() will first check for any existing *in vivo* data on bioavailability, and if not present will use calc_fbio.oral() to calculate fraction absorbed based on Caco-2 permeability. calc_fgut.oral() is available to calculate fraction surviving gut metabolism based on 1% Cl_int_. If needed, depending on how flows are described in the model, calc_hep_bioavailability() can also be used to calculate the fraction surviving first-pass hepatic metabolism.

### Calculated Chemical Parameters

Chemical-specific tissue-to-free-fraction-in-plasma partition coefficients in *httk* describe the partitioning of a chemical into various tissues (that is, equilibrium tissue:free plasma concentration ratios). These partition coefficients are estimated using a calibrated variant of the Schmitt method [60]. The specific implementation within *httk* is described in more detail by Pearce, Setzer (58). New tissues may be specified if certain composition descriptors – for example, fraction lipid – are provided. Any model in *httk* may predict partition coefficients via predict_partitioning_schmitt(), which predicts the partition coefficient for each tissue in the tissue.data table. Furthermore, users may “lump” (that is, aggregate) multiple tissue types into a single compartment with a common chemical concentration using the function lumptissues(). The specific lumping scheme (if any) used by a model is described in the corresponding model information file.

Blood to plasma ratio is another necessary parameter since most of the PBTK models in *httk* describe flows in terms of blood but concentrations in terms of tissue to plasma partition. The blood to plasma ratio helps scale available chemical fraction estimates for processes throughout the body such as partitioning, metabolism, and glomerular filtration. Although The function available_rblood2plasma() works through three options to obtain the most accurate blood to plasma concentration ratio. First, the function tries to find a species-specific measured value, otherwise it defaults to human measured values. Second, available_rblood2plasma() uses Schmitt’s method to estimate the red blood cell to plasma partition coefficient along with calc_rblood2plasma(). When insufficient chemical-specific physico-chemical data exists for the first two options the average measured value is used.

### Data Availability

There are a fixed set of chemical-specific HTTK parameters distributed with each version of *httk*. Without modifying the chem.physical_and_invitro.table these are all the chemicals that are potentially available to use in a new httk model. The *in vitro* measured values can be expanded with structure based predictions using the commands load_sipes2017() [35], load_pradeep2020() [50], and load_dawson2021() [51]. Further, only the parameters within chem.physical_and_invitro.table are available for specifying a chemical. Therefore, to develop a new HTTK model that is compatible with chemical-specific parameters available in *httk*, that model must depend on the available parameters. Currently there are the physico-chemcial properties: log_10_ membrane affinity (logMA), log_10_ hydrophobicity (logP), log_10_ water-air partition coefficient (logPwa), log_10_ Henry’s law coefficient (logHenry), log_10_ water solubility (logWSol), melting point (MP), molecular weight (MW), ionization equilibria (pKa_Accept and pKa_Donor), and chemical class. In addition, the *in vitro/in vivo* measured properties Cl_int_ (Clint), f_up_ (Funbound.plasma), Caco-2 permeability (Caco2.Pab), fraction absorbed from gut (Fabs), fraction surviving gut metabolism (Fgut), fraction surviving first-pass hepatic metabolism (Fhep), systemic oral bioavailability (Foral), and blood to plasma ratio (Rblood2plasma) are available. If a new parameter is needed (this has occurred previously) both the chem.physical_and_invitro.table and the functions for accessing it must be revised.

### Adding New Data

Existing parameterization tables are updated to incorporate new data as it becomes available. Thus, the Tables.RData file, described previously, evolves over time. Each version of *httk* potentially has a unique Tables.RData. The “Tables.Rdata.stamp” object tracks when the *httk* data tables were generated (see Table 6). The “datatables” sub-directory in the Git repository contains both the source data files and the R script “load_package_data_tables.R”. load_package_data_tables.R is used to generate the Tables.RData file from the raw data. If there is a need for a user to include additional data, they can update the load_package_data_tables.R script to re-generate the Tables.RData file with the new data. For further details, we refer the reader to the GitHub repository for *httk* (https://github.com/USEPA/CompTox-ExpoCast-httk/tree/main/datatables). To update the available data in *httk* the new Tables.RData file must be added to the “data” sub-directory. In addition, the data documentation should be updated. The “httk/R/data.R” file in *httk* contains documentation for all the data in the package. Each dataset description should specify the data object name, column names, variable descriptions, and relevant measurement units – further details on documenting R package data can be found in Section 7.1.2 of (https://r-pkgs.org/data.html) [84]. Once the documentation is complete, functions in the *roxygen2* [52, 96] package can be used to update or generate the help files accompanying data objects in *httk*.

## Model Evaluation

### Data for Model Evaluation

Model evaluation is a key part of determining whether a new HTTK model is ready for inclusion into *httk* and appropriate for use. To facilitate model evaluation several datasets are included in *httk* to aid in model evaluation. A table of concentration vs. time data is included in *httk* as a flat file along with empirical TK parameter estimates from CvTdb [38, 97]. The chem.invivo.PK.data table, mentioned in

Table 6, contains the concentration vs. time data for several dozen chemicals across multiple species (chiefly rat), often with multiple dose regimens (such as oral vs. intravenous and doses). The empirical TK estimates are included in the chem.invivo.PK.summary.data and chem.invivo.PK.aggregate.data tables, which respectively contain the estimated TK statistics from each chemical-dose regimen pair and chemical-specific parameter estimates from all available data on each chemical. Estimated TK statistics include, but are not limited to, values like C_ss_, C_max_, and AUC. Whereas chemical-specific model parameter estimates include values such as volume of distribution and elimination rate.

### Documenting Results with Vignettes

The vignettes in *httk* provide working code examples for generating figures and performing other analyses using the suite of models [52]. Vignettes are typically written in the scripting language Rmarkdown. In *httk,* we aim to provide vignettes that re-generate major figures from each HTTK-related paper. It is vital to include model assessment figures based upon model performance statistics like RMSLE. Including performance calculations in the vignette allows monitoring of changes with new version releases. Model evaluations also allow for evaluation of new functionalities. For example, if a new functionality introduces errors or inaccuracies, model evaluations within the vignettes provide a mechanism for identifying issues. A similar capability is provided by the scripts in the “tests” directory. As *httk* functions are revised and enhanced, older vignettes sometimes become obsolete. Therefore, indicating the *httk* version used when the vignette or test script was originally written is critical for accurate replication of the original analyses and figures. If an archived version of *httk* is needed, it may be obtained at: https://cran.r-project.org/src/contrib/Archive/httk/.

During the package building process, the default is to run all code chunks in every vignette. Some incompatibilities are easily caught in the vignettes during this process. This allows issues to be fixed before submitting the package to CRAN. However, some time intensive functions and code chunks may cause certain CRAN checks to fail due to time restrictions for vignette compilation. When time intensive examples are necessary, we suggest using the “eval = FALSE” option for code chunks. This allows users to provide the code for executing an analysis or figure generation but does not execute the computationally intensive code when the running the vignette for testing. The manuscript-related vignettes, which may have unevaluated code chunks because of computational requirements, are more prone to unintentional obsolescence. In addition to the manuscript-related vignettes, used for reproducing peer reviewed figures and results, we also include vignettes that serve as user guides for new *httk* users.

When a new vignette is added to the httk package it is necessary to include all the necessary data, models, and corresponding functions within the package. All relevant data should be saved as an “.RData” file in the “data” subdirectory of *httk*. Additionally, as with any other data for the package the new data should be documented using *roxygen2* [96] in “httk/R/data.R”. Any additional (non-*httk*) functions used in the vignette, which are not part of another existing R package, should be defined in the vignette Rmd or in an R script with *roxygen2* documentation [96] and placed in the “R” subdirectory of *httk*.

The default figure size for vignettes is 3 by 3 (inches). This is quite small but necessary for maintaining a package file size that meets CRAN requirements (that is, 5MB) [98]. Note, if you are using *ggplot2* [99] for figures, then to improve readability, at the default figure size, we suggest commenting out all text size specifications. The *ggplot2* package is generally good at performing the appropriate scaling when rendering figures within a vignette.

### Benchmarking *httk*

With the addition of many new models and data across versions it is not only crucial to track model performance over time, but to also track and evaluate the overall performance of *httk*. A longitudinal evaluation of the package ensures each HTTK model, and any new data included over time, is not degrading previous predictive ability. Recently, the benchmark_httk() function was added to *httk* to enable some of these evaluations and assess the impact of changes over version history – including but not limited to updates in data or code, new models, and new feature implementations – by providing predictive performance benchmarks.

For retroactive comparisons, benchmark_httk() performance statistics were obtained from previous versions of *httk* by manually re-installing each archived CRAN release of the package. Historical benchmark statistics are stored in the httk.performance table and included in the *httk* R package via the Tables.RData file. Tracking benchmark statistics allows evaluation of the predictivity of *httk* across released versions. In particular, benchmarks characterize whether the current version is performing better or worse compared to the previous versions. For adding new performance data from the latest version of *httk* to httk.performance, we refer readers to the load_package_data_tables.R in *httk* GitHub repository (https://github.com/USEPA/CompTox-ExpoCast-httk/tree/main/datatables).

Several possible benchmark are available via the benchmark_httk() function – including basic performance statistics, Monte Carlo steady state uncertainty estimations, *in vivo* statistics, and tissue partitioning coefficient checks. Table 7 provides additional details about the benchmark(s) returned from each of the checks along with brief descriptions. In addition to obtaining numeric output for each of these outputs, the ‘make.plots’ argument allows users to visualize historical performance by generating plots of performance statistics by version and including the current version. Logical arguments in the benchmark_httk() function, respective to each of the checks, allow users to control which results are reported. By default, all benchmark checks are returned. We refer readers to the benchmark_httk() help file, for further details on this function and checks.

**Table 7:**
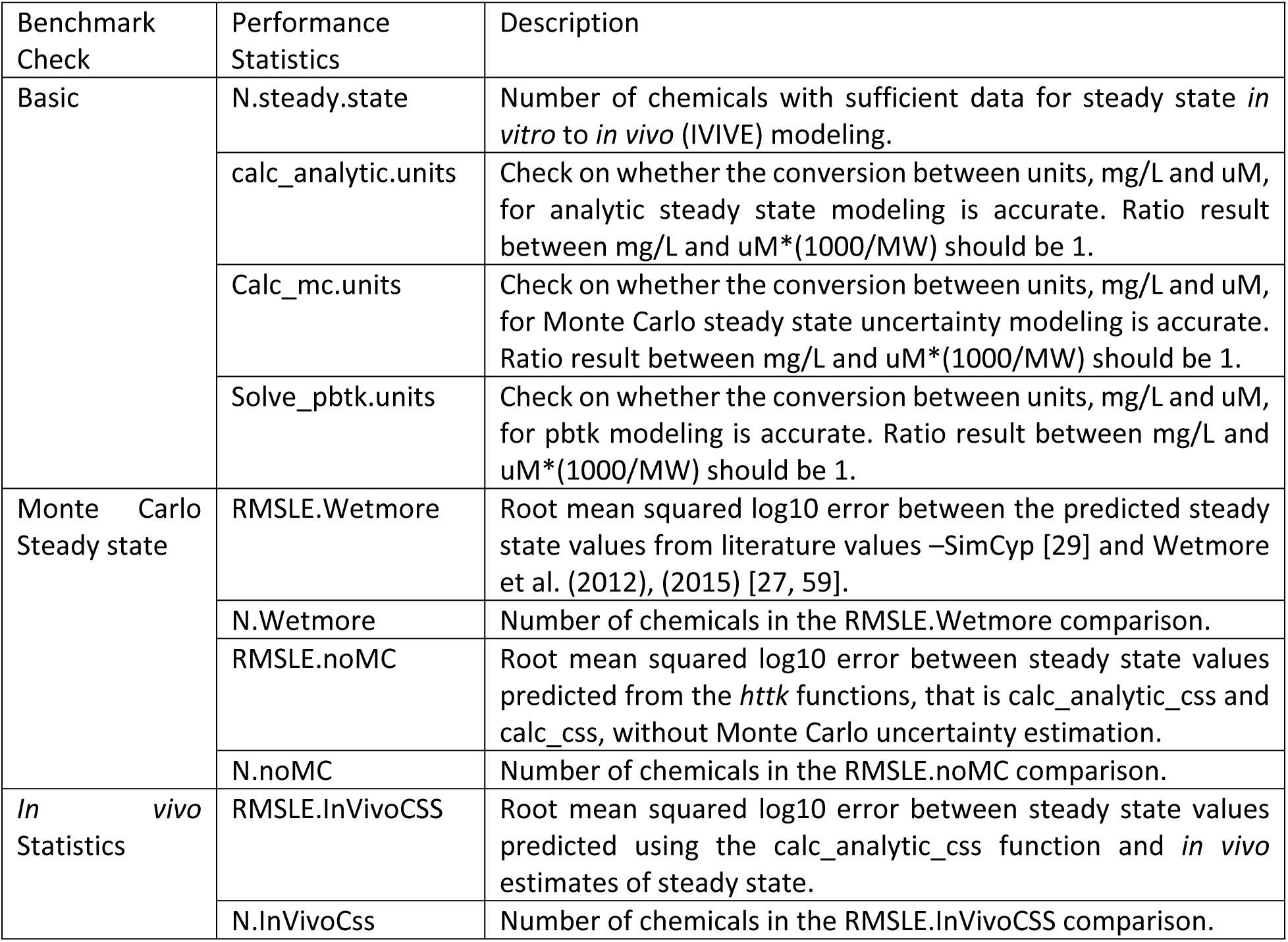
Series of performance benchmarks available through the benchmark_httk() function in httk. These performance statistics are calculated and stored in the httk.performance table for each version release of httk to track performance of the package over time.

### Generic TK Model Extensions

The previous methods sections discussed how to add a new generic HTTK model to the *httk* R package and perform basic model and package evaluations, which covers the main functionality of *httk* and the models therein. However, users may want to go beyond performing *in vitro* to *in vivo* extrapolation (IVIVE) for various exposure scenarios and quantify levels of uncertainty and variability in data and assess their impacts on model predictions. Furthermore, users may want to calculate tissue concentrations when steady state is reached (that is, metabolic equilibrium). In the *httk* package, we extend the basic functionality of HTTK models with generic functions for performing uncertainty estimation via Monte Carlo simulations and steady state calculations. Table 8 provides a list of reusable generic functions for extending the functionality of a new model.

**Table 8:** Reusable Functions for Extending the Functionality of Generic TK Models. We refer readers to the Supplemental File S2_function_guide_file.xlsx and help files for further details on these functions and other existing functions.

| Generic Functions | Basic HTTK Extension | Description |
| --- | --- | --- |
| calc_analytic_css() | Steady state Modeling | Calculates the quasi-steady state plasma concentration by calling the model-specific analytical solution when it is available for the specified model. |
| calc_css() | Steady state Modeling | This is a numerical approach to determine the quasi-steady state concentration (Css) as a function of repeated dosing. Dosing can be changed to match different conditions (default is three times a day). Also, returns the days needed to reach steady state. |
| calc_mc_tk() | Monte Carlo Simulation | Function for running a Monte Carlo simulation with a TK model. Essentially runs solve_model() for a range of parameter sets. |
| calc_mc_css() | Monte Carlo Simulation | Function for calculating Css (quasi-steady state plasma concentration) using Monte Carlo simulation of Css – typically uses the analytic solution as that is much faster than the numerical approach. |
| calc_mc_oral_equiv() | Monte Carlo Simulation | This is the “reverse dosimetry” IVIVE function that uses the Css quantiles from calc_mc_css() to estimate a daily oral equivalent dose based on quasi-steady state plasma concentration using Monte Carlo simulation. |
| calc_mc_gas_equiv() | Monte Carlo Simulation | This is a new “reverse dosimetry” IVIVE function from Breen et al. (in preparation) for daily inhalation rate that would induce a given plasma concentration at steady state. |

### Steady state Modeling

For chemical exposure simulation we are rarely interested in true “steady state” (wherein none of the state variables of the model are changing). However, we are often very interested in quasi-steady state concentrations, wherein after a sufficient time has passed for a relevant repeated discrete dosing scenario (for example, dosing three times a day), “steady state” the concentrations eventually oscillate around some average value. If the generic HTTK model you are developing is appropriate for steady state modeling then incorporating this functionality is much like the other auxiliary functions for the model described previously. The first thing that needs to be created is the model-specific analytic steady state function, namely calc_analytic_css_[MODEL]. All the necessary model equations and calculations for estimating the model-specific steady state values should be included in the calc_analytic_css_[MODEL] R function. It should be noted that the model-specific steady state function generated for a model will not be used directly. Rather, it will be used via a generic wrapper function, similar to how users interact with specific HTTK models with solve_[MODEL]() functions and the solve_model() function communicates directly with the C code for obtaining a model solution. Here, calc_analytic_css and calc_css are the generic wrapper functions that access the model-specific steady state estimating function calc_analytic_css_[MODEL](). To enable access, the corresponding generic HTTK model needs several components to be defined in the model information file – including analytic.css.func, steady.state.compartment, steady.state.units, and css.dosing.params.

Table 9 provides a brief description for each of the necessary model information components.

**Table 9:** Model information file components necessary to use steady state modeling. We refer readers to the Supplemental File S1_modelinfo_guide_file.xlsx for further details on all components.

| Model Information File Component | Description of component |
| --- | --- |
| `analytic.css.func` | Character string with the name of the model-specific steady state function, that is “`calc_analytic_css_[MODEL]`”. |
| `steady.state.compartment` | Character string for the model compartment whose steady state concentration is modeled. Typically, only one compartment. |
| `steady.state.units` | Character string providing units for the steady state modeling results returned. |
| `css.dosing.param` | Vector of character strings naming the parameters necessary to specify dosing protocols for model-specific steady state modeling, for example “`hourly.dose`”. |

After the components for the generic HTTK model of interest are included in the model information file, the calc_analytic_css() and calc_css() functions can perform steady state modeling when the argument “model” indicates the model of interest. For further details on constructing the calc_analytic_css_[MODEL] function and updating the model information file, we refer the reader to the help files and vignette in the *httk* package.

### Monte Carlo Simulation

Within *httk* uncertainty and variability in the model parameters can be propagated to the model output with Monte Carlo (MC) simulation [19]. In MC simulations model parameters are sampled from user-specified distributions and the model is evaluated for each set of parameters. The parameters may be sampled from different distributions depending on the user’s preferences. For physiological parameters in particular, the user may choose to apply *httk*’s built-in population physiology sampler known as HTTK-Pop [40]. Here, we describe the updates necessary for both types of MC simulation together and denote which updates are specific to HTTK-Pop. It should be noted that use of HTTK-Pop is not required, and HTTK-Pop should not be used for non-human populations. Extending the basic functionality of a generic HTTK model to include MC simulation capability follows a process similar to the steady state modeling extension described above.

To understand the necessary components for enabling the MC simulations capability with a new model, it is useful to understand *httk’s* built-in wrapper functions that perform MC. The built-in wrapper functions for MC simulation, such as calc_mc_css() and calc_mc_oral_equiv(), were described previously in Table 8. Both calc_mc_css() and calc_mc_oral_equiv() require the analytical steady state calculation function, calc_analytic_css(), to already be incorporated – see the Steady state Modeling section for further details. Figure 5 provides an overall schematic of the MC simulation process, and highlights some of the model-specific pieces needed when extending an HTTK model.

**Figure 5:**
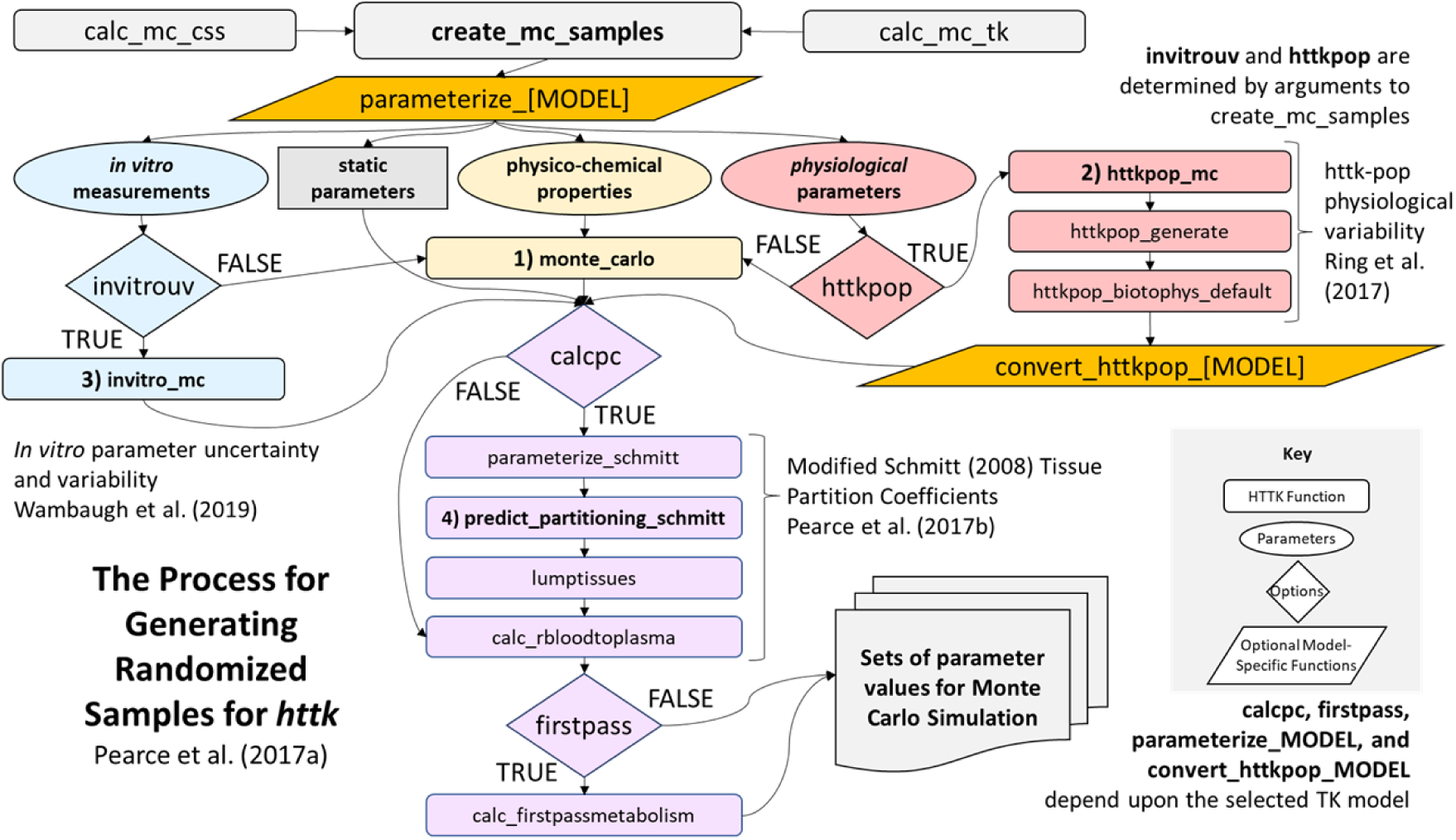
Schematic of the overall Monte Carlo (MC) simulation process for propagating uncertainty and variability in httk (Figure adapted from [41]). The blue box, create_mc_samples(), denotes the built-in wrapper function coordinating model-specific information from the modelinfo file and user specified arguments to generate MC samples. The yellow boxes, parameterize_[MODEL]() and convert_httkpop_[MODEL](), are model-specific functions that may be necessary for integrating MC sampling. The red boxes, calcpc and firstpass, denote model-specific logical arguments specified in the modelinfo file for a given HTTK model. Note, this figure is not meant to provide an exhaustive list of all possible components that may be necesssary to enable MC simulation capabilities for a generic HTTK model.

Both wrapper functions, use create_mc_samples() to generate a table of MC-sampled model parameters. Users may also call create_mc_samples() directly. To incorporate a new model into the existing built-in MC wrapper, functions require the user to define several components in the model information file. Table 10 provides a list of MC simulation components necessary for the model information file with a brief description of each. It should be noted that some components are HTTK-Pop specific, and some are optional based on additional functionality that may be relevant for a given model.

**Table 10:**
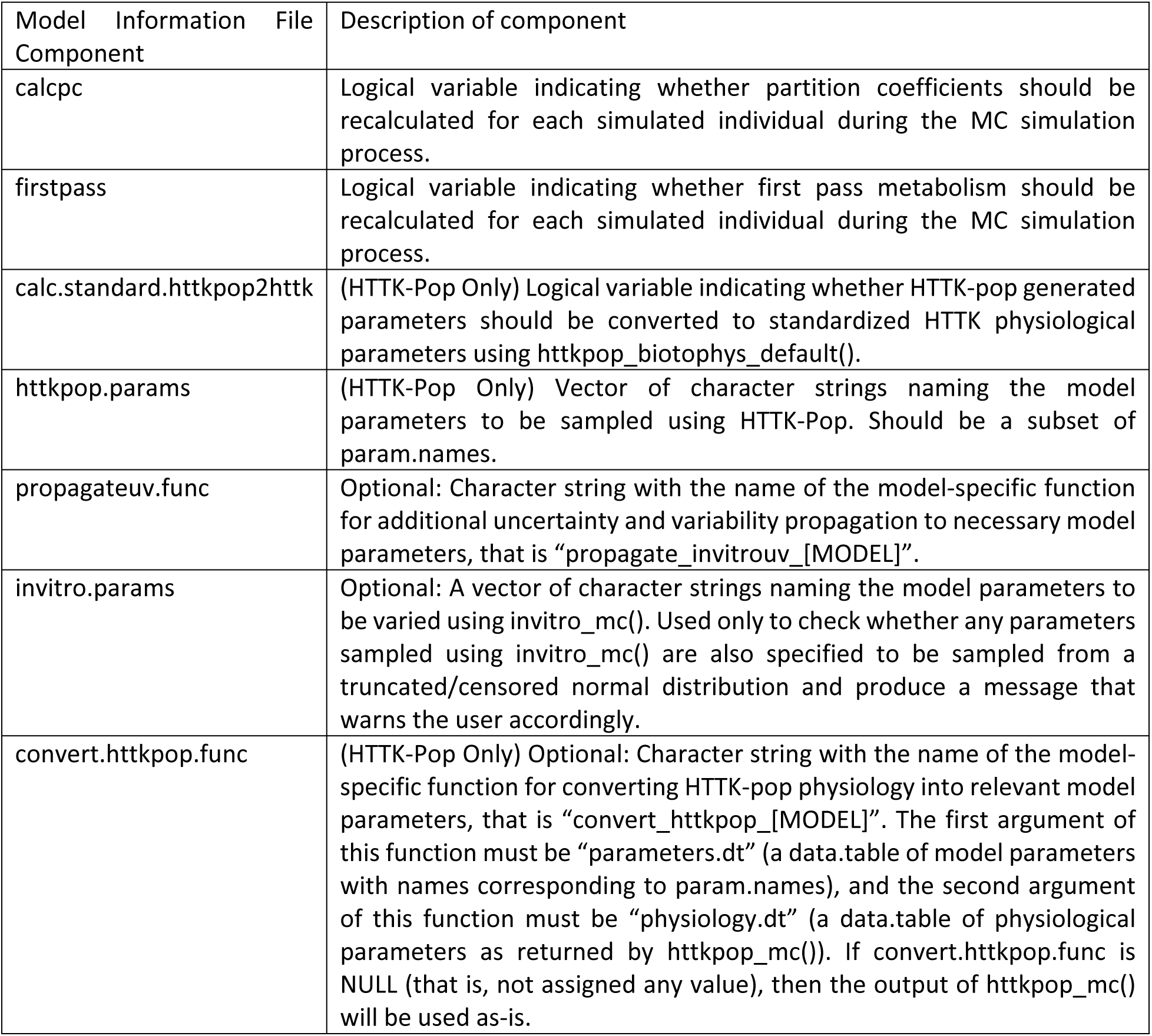
Model information file components necessary to use Monte Carlo simulation. This list does not include the elements parameterize.func and param.names, which are assumed to be part of the existing basic components in the model information file – see Table 4. We refer readers to the Supplemental File S1_modelinfo_guide_file.xlsx for further details on all components.

The create_mc_sample() function uses the components listed in Table 10 at various in Figure 5. Here, we will briefly describe how create_mc_sample() works and explain when each component is required. First, create_mc_samples() generates the default set of parameters for the model, using the parameterize_[MODEL] function defined under “parameterize.func”, see the Basic Generation and Integration of a New HTTK Model section for further details. Next, MC simulation of parameters is done in four steps, some of which may be optional depending upon relevant functionality. These steps include drawing samples for relevant parameters from truncated/censored distributions, sampling physiological parameters using HTTK-Pop, sampling chemical-specific *in vitro* parameters, and recalculating parameter values that depend upon other parameters.

Step One draws samples for any parameters listed in the “censored.params” and/or “vary.params” arguments of create_mc_sample() from censored and/or truncated normal distributions, respectively. Nothing needs to be defined in the modelinfo file to complete this step. However, if the components “httkpop.params” or “invitro.params” are defined (see Table 10), they are used to check whether a parameter sampled as part of this step will be overwritten in Steps Two or Three of MC simulation and will produce a warning message for the user.

Step Two, HTTK-Pop sampling, only occurs if the “httkpop” and “species” arguments for create_mc_samples() are set to TRUE and “human”, respectively. The httkpop_mc() function will sample physiological parameters from distributions based on the U.S. population as surveyed by CDC NHANES [41, 100–102]. If a modeler desires parameters returned by httkpop_generate() to be converted to standardized units expected by the existing HTTK models (that is, directing httkpop_mc() to run httkpop_biotophys_default()), then the “calc.standard.httkpop2httk” in the modelinfo file should be TRUE. For further details on the httkpop_biotophys_default() function we refer readers to the *httk* help file. The resulting sampled parameter table will be returned to create_mc_samples(). When additional conversions or calculations on the parameters output by httkpop_mc() are necessary, the user may optionally define a model-specific function, namely convert_httkpop_[MODEL](), and provide the name to the “convert.httkpop.func” component in the modelinfo file. If “convert.httkpop.func” is defined, then create_mc_samples() calls the specified function. Otherwise, the output of httkpop_mc() will be used as-is. After httkpop_mc() and (if used) convert_httkpop_[MODEL]() are called, parameters in the resulting table will be retained only if they appear in the parameter list returned by the parameterize_[MODEL] function.

Step Three, if the “invitrouv” argument for create_mc_samples() is TRUE, then the invitro_mc() function performs Monte Carlo sampling for *in vitro* chemical-specific parameters – such as Cl_int_ (Clint), fraction unbound in plasma (Funbound.plasma), and Caco2 permeability (Caco2.Pab). Nothing needs to be specified in the modelinfo file for this step.

Step Four, the create_mc_samples() function recalculates parameter values that depend on the values of other parameters – particularly values of *in vitro* chemical-specific parameters sampled in Step Three. Modelers may control this process by specifying the “calcpc” and “firstpass” components, both of which are logical – that is TRUE/FALSE, in the modelinfo file. If “calcpc” is TRUE, then the tissue:plasma partition coefficients are recalculated for each simulated individual using the individual’s value for fraction unbound in plasma (which may be varied in Step Three) and other relevant parameters. If “firstpass” is TRUE, then the partition coefficients are recalculated, and first-pass hepatic bioavailability is also recalculated for each simulated individual. For additional model-specific uncertainty/variability propagation that may be necessary, a modeler may define a model-specific function, namely propagate_invitrouv_[MODEL](). For example, propagate_invitrouv_1compartment() computes “Vdist” (volume of distribution) and “kelim” (elimination rate), in addition to the standard parameters, using the other sampled parameter values. The first argument for the propagate_invitrouv_[MODEL]() function should be “parameters.dt”, which accepts a data.table of model parameters. Other arguments available for the function are those that can be passed into the “propagate.invitrouv.arg.list” argument of create_mc_samples(). A data.table of model parameters is the expected output for this function. If a modeler defines a model-specific uncertainty/variability propagation function, then it should be specified in the modelinfo file component “propagateuv.func”.

After the creating the model-specific functions and updating the relevant components in the model information file, for the generic HTTK model of interest, the create_mc_samples() function can generate the necessary samples for MC simulations with the argument “model” indicating the model of interest. We refer the reader to the *httk* help files and vignettes for further details on constructing functions and updating the model information files for MC simulations.

## Acknowledgements

The authors thank Drs. Dustin Kapraun, Kelsey Vitense, and Todd Zurlinden for their constructive feedback during the U.S. EPA internal reviews of the manuscript. We appreciate Dr. R. Woodrow Setzer’s insights into dynamic modeling in R and statistical analysis of PBTK models. We appreciate programming support from Mohideen Marikar.

## Disclaimer

The views expressed in this publication are those of the authors and do not necessarily represent the views or policies of the U.S. EPA. Reference to commercial products or services does not constitute endorsement.

## Data Availability Statement

R package “httk”, including R scripts, data files, and C code, is freely distributed via the Comprehensive R Archive Network (CRAN – https://cran.r-project.org/). The “httk” codebase is provided in a public GitHub repository (https://github.com/USEPA/CompTox-ExpoCast-httk). A change log describing revisions is available at: https://cran.r-project.org/web/packages/httk/news/news.html. HTTK models can be found at: https://github.com/USEPA/CompTox-ExpoCast-httk/tree/main/models.

## Supplemental Material

- model_gas_pbtk.model – MCSim [85] computer language description of Linakis et al.[7] model
- model_gas_pbtk-raw.c – C computer language [103] description of the [7] model as initially generated by MCSim
- model_gas_pbtk.c – C computer language description of the [7] model as modified for integration into httk
- modelinfo_gas_pbtk.R – R computer language [104] documentation of the [7] model for “model aware” httk functions
- parameterize_gas_pbtk.R – R computer language function for generating chemical-specific parameters for the [7] model
- solve_gas_pbtk.R – R computer language function for simulating the [7] model
- Vh_Linakis2020.Rmd – Rmarkdown vignette for generating the figures from [7]
- S1_modelinfo_guide_file.xlsx – Excel file containing brief descriptions of modelinfo file components and organizing components by their relevant role in adding/extending a generic HTTK model in *httk.* Current as of April 2024.
- S2_function_guide_file.xlsx – Excel file containing brief descriptions of functions within the R package, which modelers may want to use when integrating/extending their new generic HTTK model into *httk.* Current as of April 2024.

